# Lifetime exposure to known and emerging groundwater contaminants significantly alters poultry microbiome and metabolome

**DOI:** 10.1101/2025.07.12.664519

**Authors:** Chamia C. Chatman, Elena G. Olson, Steven C. Ricke, Erica L.-W. Majumder

## Abstract

The exposome encompasses all lifetime environmental exposures affecting health. Its complexity and high data dimensionality make it challenging to link specific exposure combinations to adverse health outcomes. Establishing relevant exposome criteria is key to addressing current knowledge gaps. This study evaluated contaminant levels in Wisconsin groundwater and their effects on host health. We focused on three co-occurring chemicals that were detected at concentrations exceeding groundwater standards, nitrate, atrazine and imidacloprid, and the emerging contaminant, microplastics. In this study, broilers were exposed to a low dose chemical mixture (35,000 ppb nitrate + 1.7 ppb atrazine + 0.58 ppb imidacloprid) and high dose chemical mixture (100,000 ppb nitrate + 3,000 ppb atrazine + 3,000 ppb imidacloprid) or polyethylene microplastics (PE MPs) for 49 days. We observed that both ternary mixtures and PE fiber MPs significantly altered the cecal microbiomes as determined by the enrichment of genera, *Fournierella*, *Ruminococcus* and an unclassified genus in the family *Coriobacteriaceae*. In addition, +PE fiber presence dysregulated metabolic pathways associated with bile acid biosynthesis and lipid metabolism. Similarly, perturbations to cecal microbial activity for both ternary chemical mixtures were confirmed via modulation of six metabolites including methylisopelletierine which had a higher total ion intensity than the control group. Interestingly, there were no detectable pathological effects to either the +PE fiber or ternary mixture treatment groups. Overall, the data presented here demonstrates that low doses of environmental contaminants are sufficient to dysregulate cecal taxonomic composition and microbial activity without inducing detectable pathological effects.

**Importance:** We found that exposure to mixtures of environmental toxins caused gut dysbiosis observed by changes to the chicken cecal microbiome and metabolome. This highlights the importance of conducting such studies with environmentally relevant mixtures of contaminants at detected concentrations to understand the actual risks associated with exposures like drinking contaminated groundwater over a long period of time. Our findings demonstrate that gut microbial metabolites, now known to be key regulators and signaling molecules in a wide range of host health issues, are the source of the negative health outcomes; superseding cell death or pathological damage that are caused by acute exposures. These changes have implications for predicting negative long-term chronic health outcomes.

## 1. Introduction

There has been an observed increase in the presence of agricultural chemicals in drinking water (1). However, the potential risks or health effects associated with chemical mixtures detected in groundwater remain poorly understood due to the lack of chemical mixture studies performed at environmentally relevant concentrations (2).

Xenobiotics are defined as any exogenous substances or microorganisms that enter a host (3). Xenobiotics detected in groundwater can be categorized as known contaminants (i.e., pesticides) or emerging contaminants (i.e., microplastics). Humans and livestock are primarily exposed to agricultural chemicals or microplastics via ingestion (3–6). However, inhalation and dermal contact are also sources of exposure for microplastics (5).

A central role of the gut microbiota is biotransformation of ingested molecules to bioactive compounds which directly influence mucosal integrity, immune function and energy metabolism (3). Xenobiotic metabolism is accomplished via phase I or phase II metabolism within the small intestine and liver (7, 8). Despite the ability of the gut microbiota to biotransform xenobiotics, agricultural chemicals and microplastics have been demonstrated to perturb the gut microbiome. For example, Yu et al. (2023) observed that turbot, a fish species native to Europe, exposed to nitrate experienced significant decreases in *Firmicutes* and *Cyanobacteria* in a dose-dependent manner (9). In frogs exposed to chronic levels of atrazine (50 µg/L to 500 µg/L) it was reported that there was significant modulation of *Fusobacteria* and *U114* (10). In addition, Huang et al. (2022) demonstrated via liquid-chromatography mass spectrometry that microbial activity was altered in the intestine of <u>Pelophylax</u> *nigromaculatus* larvae following chronic atrazine exposure as evidenced by the significant dysregulation of pathways associated with purine metabolism and amino acid biosynthesis (10). Interestingly, honeybees exposed to imidacloprid did not exhibit signs of perturbed gut microbiomes, but increased mortality occurred (11). However, zebrafish exposed to 1000 μg/L imidacloprid (1,000 ppb imidacloprid) did elicit a significant impact on gut microbial composition (12). This highlights the variability in species sensitivity to agricultural chemicals, including nitrate, imidacloprid, and atrazine.

Microplastics (MPs) may enter terrestrial environments as either primary or secondary microplastics. Primary MPs are polymers that are designed to be used in commercial products like cosmetic products, personal care products, pharmaceuticals and detergents (13, 14). Secondary MPs represent larger pieces of plastic which have been degraded into smaller polymers via physical, chemical or biological interactions in the environment (14). Thus, resulting in the formation of microplastics which are polymers less than five mm in diameter (14). In recent years, MPs have been detected in human placenta (15, 16), testes (17, 18) and other human tissues (13) which has exacerbated public health concerns. Research has also indicated that MP characteristics such as polymer type, shape and size have varied biological impacts (4–7). Humans may become exposed to MPs primarily via ingestion; however, inhalation and dermal contact are also sources of exposure (8). Livestock such as chickens may also be exposed to MPs via ingestion (19).

Here, we evaluated effects of known contaminants, a mixture of nitrate, imidacloprid and atrazine, and the emerging contaminant, polyethylene fiber microplastics (PE Fiber MPs) using broiler chickens as an *in vivo* model system.

Broilers are the most consumed meat product in the United States of America (USA) (20), however; there is lack of knowledge on the effects of known contaminants (i.e., pesticides) or emerging contaminants (i.e., microplastics) on their long-term health. Additionally, the ceca are the primary site for microbial activity in the poultry gastrointestinal tract (21) and thus an ideal environment to explore xenobiotic metabolism. Furthermore, integration of 16S rRNA sequencing and untargeted metabolomics provides a multi-omics approach enabling a deeper understanding of the microbiome’s functional potential, especially as it pertains to interactions with environmental contaminants (22). Nitrate, imidacloprid and atrazine were selected as the agricultural chemical mixture due to their frequency of detection and concentrations exceeding enforcement standards in the 2021 Department of Agriculture, Trade and Consumer Protection (DATCP) Targeted Sampling Report (23). In addition, PE MPs were incorporated as the emerging contaminant of interest based on our previous research which indicated that polyethylene (PE) fiber MPs alter the taxonomic composition of cecal microbiomes within *in vitro* cecal mesocosms (24). Given the health effects of each of these agricultural chemicals singularly and previous evidence that PE fiber alter the cecal microbiome (24), we hypothesized that exposure to the chemical mixtures and PE fiber MPs would alter the broiler cecal microbiome and subsequently impact cecal microbial activity via modulation of key metabolic pathways *in vivo*. We observed that PE fiber MPs significantly altered poultry cecal microbiome composition and microbial activity. Interestingly, the +PE Fiber group was enriched in *Fournierella,* but depleted in *Synergistes* indicating potential beneficial impacts to PE fiber presence, although untargeted metabolomics results highlighted significant dysregulation of metabolic pathways in the +PE Fiber treatment group which are associated with increased incidence of metabolic disorders. Similarly, exposure to a ternary mixture of nitrate, atrazine and imidacloprid, even at environmentally relevant concentrations, significantly altered metabolic pathways such as metabolism of xenobiotics by cytochrome P450, pyruvate metabolism, and thiamine metabolism.

Overall, we observed significant impacts to cecal microbial taxonomic composition and microbial activity due to the presence of these environmental contaminants.

## 2. METHODS

### 2.1. Animal Husbandry and Experimental Design

This experiment was conducted with the consent of the Institutional Care and Use Committee at the University of Wisconsin-Madison (protocol: A006804). Aviagen Ross 308 Broiler chickens (mixed sex) were purchased from Welp Hatchery in Bancroft, Iowa. Broilers were housed at the Poultry Research Facility at the University of Wisconsin-Madison. Birds were randomly assigned to a treatment group and immediately placed into the designated floor pen. The broilers (N=100) were allowed to acclimate to the facility for 7 days. Water and commercial poultry diet (Agrimaster 21% Meat Producer Poultry Feed Crumbles; Blain # 334839, Mfr # 3931) were fed *ad libitum* (Table S1-S2). The Agrimaster diet is designed to be fed to poultry during the starter, grower, and finisher phase of broiler production.

### 2.2. Preparation of polyethylene microplastics

Following methods by Cole (2016) (25), low density polyethylene (LDPE) fibers (GoodFellow, LS554234) were cut to 50 µm using a cryotome (Leica biosystem 1950 cryotome). Polyethylene fiber sizes were subsequently verified using scanning electron microscopy (Zeiss GeminiSEM 450) (26). Polyethylene fibers (250 mg/kg poultry basal diet) were then added into to the poultry basal diet weekly to account for changes in feed consumption during the starter, grower and finisher phase of broiler production. For this study, the microplastic concentration was based on the median concentration of microplastics detected in a terrestrial environment in the USA (27). A similar concentration was also used for an *in vivo* MP toxicity study (28).

### 2.3. Selection and Preparation of Groundwater Contaminant Chemicals

We selected a mixture of nitrate, atrazine and imidacloprid based on their frequency of detection and concentrations exceeding enforcement standards in the 2021 Department of Agriculture, Trade and Consumer Protection (DATCP) Targeted Sampling Report (23). Chemicals used in this study were atrazine (ASTATACH, Lot: P151-04287), imidacloprid (Thermo Scientific, Lot: A0450203), and nitrate (AccuStandard, Lot: 222105041). Atrazine was first diluted in dimethyl sulfoxide (DMSO) then final dilutions were done in autoclaved ultrapure water. Nitrate and imidacloprid dilutions were done in autoclaved ultrapure water.

In our previous study, nitrate, atrazine and imidacloprid were selected due to their frequency of detection in groundwater wells and chemical concentrations for each exceeding either an enforcement standard or preventative action limit (29). One of the three treatment groups represented chemical concentrations analogous to those detected in the 2021 DATCP Targeted Sampling Report. We then used *in silico* and *in vitro* methodologies to further evaluate the DATCP report and elucidate if potential chemical-biological interactions would be present between the chemical mixtures and our selected models, Caco-2 cells and poultry cecal microbiota. We concluded that the poultry cecal microbiota exposed to the ternary mixtures had a decline in growth rate (29). Whereas, Caco-2 cells exposed to various two-chemical combinations of nitrate, atrazine and imidacloprid did elicit a significant decrease in cell viability, but not the ternary mixtures (29). For this *in vivo* trial, the low dose chemical mixture represents chemical concentrations like concentrations detected in the 2021 DATCP Targeted Sampling Report (23) (Table 1). The high dose mixture was based on preliminary exposure studies where it was observed that two-chemical combinations of each chemical (i.e., nitrate + atrazine, nitrate + imidacloprid and atrazine + imidacloprid) at 3,000 ppb resulted in growth inhibition to poultry cecal organisms *in vitro* (*29*). The concentration of nitrate was set higher than detected levels and greater than the preventative action limit for nitrate (30). Furthermore, atrazine and imidacloprid concentrations used for the high dose chemical mixture exceeded the preventative action limits for each chemical (30).

**Table 1.** Composition of agricultural chemical mixtures used in this study.

|  | **Nitrate**<br>**Concentration (ppb)** | **Atrazine**<br>**Concentration (ppb)** | **Imidacloprid**<br>**Concentration (ppb)** |
| --- | --- | --- | --- |
| Low dose | 35,000 | 1.7 | 0.58 |
| High dose | 100,000 | 3,000 | 3,000 |

### 2.4. 49-day MP exposure study in broilers

Following the acclimation period, broilers were randomly assigned to one of four treatment groups. The treatment groups are as follows: untreated control group (n=17), Polyethylene fiber treatment group (+PE Fiber; n=20), low dose mixture (n=22), and high dose mixture (n=19). The treated feed was prepared weekly by pouring each treatment into a designated feed bin while gently mixing the feed. On day 49, all broiler chickens were humanely euthanized with carbon dioxide asphyxiation. Ceca from each bird was separated at the ileal-cecal junction with sterile scalpel blades and disposable forceps. The cecal digesta was divided into two aliquots and flash-frozen with liquid nitrogen to be further processed for 16S rRNA sequencing and untargeted metabolomics analyses. A section of the spleen, liver, kidney, duodenum, pancreas and cecum from each broiler were also removed from each broiler using sterile scalpel blades and disposable forceps for hematoxylin and eosin (H & E) staining.

### 2.5. Evaluation of growth parameters

Individual bird weights were recorded on days 0, 7, 14, 28, 35, 42 and 49.

Broilers’ individual weights were subsequently averaged to account for total body weight gain per treatment (accounting for mortalities). Body weight gain (BWG) was defined as the ratio of total body weight gain and the number of days on trial. To monitor feed consumption, the feeders were also weighed per pen on days 0, 7, 14, 28, 35, 42 and 49. Mortalities were not included when calculating feed conversion ratios due to final carcass weights not being recorded. Feed conversion ratio (FCR) is the ratio of the total feed intake (per pen) and total weight gain (per pen). Feed intake (per pen) is defined here as the difference in weight of feed bins pre- and one- week post-feeding for each treatment group. One-way analysis of variance (ANOVA) statical analysis was performed on BWG parameters in GraphPad Prism (version 10.4.0) (Brown-Forsythe and Welch ANOVA test; P-value ≤ 0.05). To further compare mean BWG across groups, a Dunnett’s multiple comparison test was conducted (P-value ≤ 0.05).

Descriptive statistics were performed in GraphPad Prism (version 10.4.0) to compare mean feed conversion ratio across treatment groups. Statistical analysis was then performed for feed intake (FI) data in R (version 4.4.0) (Linear Mixed Effects Model; P- value ≤ 0.05).

### 2.6. Histological evaluation

All the tissues were evaluated blindly and scored by American College of Veterinary Pathologist (ACVP) board-certified Veterinarian Pathologist at the University of Wisconsin-Madison’s Comparative Pathology Laboratory using standard methods (31–33). In brief, tissues were harvested from the same anatomical location from all the animals (N= 78). Tissues evaluated included liver, spleen, kidney, duodenum, pancreas, ceca, and colon. All tissues were collected and fixed in 10% buffered formalin for 24 hours and transferred after to 70% ethyl alcohol until processing. Formalin-fixed tissue sections were processed by a Sakura Tissue-TEK VIP, and paraffin-embedded before sectioning onto a glass slide. Tissue sections (5µm) were observed using an optical microscope (Olympus Bx 43, Software CellSens Standard). The table scoring system used a four-point scale (0-none, 1-minimal, 2-mild, 3- moderate, and 4-severe changes). Tissues sections were evaluated for the following perturbations. Livers were assessed for vascular changes included peliosis hepatis-like lesions, apoptosis, vacuolar changes, and hepatic lipidosis; spleens for lymphoid depletion; kidneys for renal tubular hydropic/vacuolar degeneration and formation of vacuoles; small and large intestines for changes of enteric and enteric epithelial architecture; and pancreas for exocrine and endocrine pancreas.

### 2.7. Library Preparation and 16S rRNA Gene Amplicon Community Sequencing

DNA extraction was performed using the standard protocol for the DNeasy Blood and Tissue kit (Qiagen, Cat 69506). Cecal samples from each bird were collected and stored in -80°C until genomic DNA extractions were performed (N= 78; control = 17, +PE fiber = 20, Low dose chemical mixture = 22 and High dose chemical mixture = 19). A 0.5 mL aliquot from each bird was used for gDNA extractions. Samples were centrifuged for 5 min x 14,000 x g and the supernatant was subsequently discarded.

Total genomic DNA quality and concentration were verified using an Infinite 200Pro spectrophotometer (Tecan Nano Quant Plate™). The DNA extracts were diluted to 10 ng/µL in Buffer AE and stored at -80°C until further analysis.

To initiate amplicon library preparation for microbiome sequencing, DNA extracts were PCR-amplified with primers targeting the V4 region of the 16S rRNA gene. The primers were dual-indexed primers designed using high-fidelity polymerase, PhiX, according to the protocol by Kozich et al (2013)(34). Gel electrophoresis was performed to verify amplified PCR products. The amplified PCR products were then normalized to 20 µL using a SequalPrep Normalization kit (Life Technologies, Carlsbad, CA, United States). Aliquots of 5 µL from each normalized sample were subsequently pooled into the final library. Final concentrations were verified using a KAPA library quantification kit (Kapa Biosystems, Wilmington, MA, United States) and a Qubit 2.0 Fluorometer (Invitrogen, Waltham, MA, United States). Next, the final library was diluted to 20 nM with HT1 buffer and PhiX control v3 (20%, v/v). Then 600 µL of the final library was loaded onto a MiSeq v2 (500 cycles) reagent cartridge (Illumina, San Diego, CA, United States) to begin the sequencing run.

### 2.8. Microbiome community analyses

The raw Illumina amplicon sequence data were uploaded to BaseSpace website (Illumina, San Diego, CA, United States) to assess sequence run quality and completion. Quantitative Insights Into Microbial Ecology (QIIME) 2 (version 2024.5) (35) was utilized to perform microbiome bioinformatics. The demultiplexed data were downloaded from the Illumina Basespace website and uploaded to QIIME2 using Casava 1.8 paired-end demultiplexed format (via QIIME import tools). The DADA2 (36) data (via q2-dada2) was subsequently used to quality-filter using the chimera consensus pipeline. Microbiome samples used for comparing control versus +PE Fiber treatment groups were filtered to a sampling depth of 3,002 which retained 84, 056 features (16.70%) in 77.78% of samples (N= 28). Microbiome samples comparing control, low dose mixture and high dose mixture were filtered to a sampling depth of 1,143 which retained 49,149 features (8.46%) in 74.14% of samples (N= 43). Alpha rarefaction plots were used to confirm the sampling depth. The phylogenetic tree (via q2-phylogeny) was generated by aligning the amplicon sequence variants with mafft (37) (via q2-alignment) and fastree2 (38). Taxonomy was subsequently assigned to the ASVs in the feature table using (via q2-classifier sklearn) (confidence limit of 97%) (39). The classifier was trained using the SILVA 138 99% operational taxonomic unit reference sequences (40).

To identify differentially abundant features across treatment groups, Analysis of Compositions of Microbiomes with Bias Correction (ANCOM-BC) was used. ANCOM-BC is a linear regression model that corrects for bias by accounting for differential sampling fractions across samples (41). A pairwise comparison using Kruskal-Wallis was conducted for α-diversity metrics: Shannon’s Diversity Index, Observed Features, Faith’s Phylogenetic Diversity, and Pielou’s Evenness (P-value ≤ 0.05;Q-value ≤0.05)(42). Main effects and interactions of variables were assessed with analysis of variance (ANOVA) for α-diversity metrics. Treatment effects based on β-diversity metrics (i.e., Jaccard distance, Bray-Curtis distance, unweighted UniFrac distance, and weighted UniFrac distance) (via q2-composition) were assessed with non-parametric multivariate analysis of variance (PERMANOVA) (P-value ≤ 0.05; Q-value ≤0.05)(43, 44).

### 2.9. Metabolite Extraction

Prior to metabolite extraction, ∼1 g of cecal digesta was diluted in 2 mL of autoclaved water. An aliquot (0.3 mL) of the diluted cecal digesta was used to determine protein concentration by a Bradford assay (45). Metabolite extraction was performed using 0.5 mL of diluted cecal digesta from each broiler. Cells were lysed by three freeze-thaw and sonication cycles, which consisted of thawing at room temperature for 10 min followed by a 30-s sonication on ice in a water bath sonicator (Branson 2800). A 1 mL aliquot of 2:2:1 (vol/vol) mixture of ≥99.9% purity acetonitrile (Fisher Scientific; Cat.A998-4) ≥99.8% purity methanol (Fisher Scientific; Cat. A412-4), and water was added to cell extracts and sonicated for 30 s and stored at −20°C overnight to allow cellular debris and protein to precipitate. The next day, samples were centrifuged for 15 min at 20,784 × g at 4°C. The supernatant was transferred to new microcentrifuge tubes and dried for 6 h on a SpeedVac Concentrator (Thermo Scientific Savant DNA 120). Dried samples were then reconstituted in acetonitrile:water (1:1 vol/vol) solvent mixture based on the normalized protein concentration of all samples, where the highest protein concentration sample had 100 μL of resuspension volume.

Samples were vortexed for 30 s, sonicated on ice for 10 min, and centrifuged for 15 min at 13,000 rpm at 4°C to remove any residual debris. The metabolite extracts were transferred into 0.3 mL high performance liquid chromatography (HPLC) autosampler vials with inserts (VWR International LLC, Cat. 9532S-1CP-RS) and stored in −80°C until analysis.

### 2.10. Untargeted metabolomics

Using a protocol by Chatman et al. (2024) (24) untargeted metabolomics was conducted. Metabolite extracts were analyzed with ultra-high-performance liquid orbitrap chromatography mass spectrometry (UHPLC-MS) (Thermo Scientific Orbitrap Exploris 240 mass spectrometer). A Kinetex Core-Shell 100 Å column C-18 column (1 × 150 mm, 1.7 μm, Phenomenex) was used for separation of metabolites. Sample injection volume was 3 μL at a flow rate of 0.250 mL/min. Mobile phase A was composed of water with 0.1% formic acid, and mobile phase B was 0.1% acetonitrile.

The gradient began with 5% B from 0 to 3 min, then an increase and hold to 95% B until 18 min, followed by a decrease to 5% B at 18.50 min and held at 5% B until 22 min.

Data-dependent acquisition was used for the tandem MS workflow in positive ion mode. Twenty-eight of the extracts were run successfully before an UPLC malfunction; however, due to our practice of randomizing samples in the run order and including QA and QC samples, there were sufficient replicates from each group to proceed with metabolomics data analysis.

### 2.11. Untargeted metabolomics data analysis

MetaboAnalyst 6.0 was used for statistical analysis and functional analysis of the MS1 data as described in Pang et al. (2022) (46). The raw files were converted to mzML files and centroided (MS-1, orbitrap, positive mode) using ProteoWizard version 3.0 (47). Univariate analyses were performed to investigate statistical differences in metabolite features between treatment groups. Principal component analysis provided insight into similarities and differences among the treatment group. Analysis of variance analysis (ANOVA) was conducted to investigate potential differences in metabolite features between each treatment group (Tukey’s post-hoc test; P≤ 0.05). Hierarchical clustering was also conducted to evaluate the top 25 features across treatment groups (distance measure: Euclidean, clustering method: Ward; P-value = 0.05). Functional analysis (P-value = 0.05, KEGG *Gallus gallus* library) was conducted to assess pathway-level changes to the cecal metabolome. Feature annotation of metabolites was first carried out using MetaboAnalyst 6.0, which matches MS1 data compounds in Human Metabolome Database. Functional analysis was subsequently conducted to assess pathway-level changes to the cecal metabolome. Compound Discoverer (Version 3.3, Thermo Fisher Scientific) was used for statistical, identification and functional analyses of raw MS/MS data. Raw MS/MS files in Compound Discoverer were compared against mzCloud, Metobolika, and ChemSpider databases.

## 3. RESULTS

### 3.1. Environmental contaminants did impact growth performance but did not induce pathological damage

To assess the impact of PE fiber MPs on broiler growth performance, average body weight gain (BWG), feed intake (FI) and feed conversion ratio (FCR) were calculated per pen (Table 2). Overall, there was no significant change to BWG from D0 to D49 when comparing control and +PE fiber MPs treatment group (Table 2). However, we observed that BWG for the +PE fiber treatment group was significantly lower at D28 to 35 compared to the control (P-value ≤ 0.05) (Table 2). Analysis of the mean FCR over the treatment period (49 days) indicated that the +PE Fiber treatment group had a lower mean FCR (Table 2). Analysis of FI was also performed to determine if exposure to agricultural contaminants negatively impacted broilers’ feed consumption. The +PE fiber treatment group exhibited significantly different intake patterns from control birds (Figure S1; P < 0.001). Though FI patterns appear similar, the presence of PE Fiber in the poultry diet increased FI at later growth stages, particularly after D28 (Figure S1; P < 0.001).

**Table 2.** Summary of body weight gain (BWG) and feed conversion ratio (FCR) results comparing control, and +PE fiber treatment groups.

|  |  |  |  |  |
| --- | --- | --- | --- | --- |
| **BWG (kg/bird/week)** | **Control Mean (n = 17)** | **+PE Fiber mean (n=20)** | **P-value<sup>a</sup>** | **SEM** |
| *D0 to 7* | *0.00* | *0.00* | *0.16* | *0.00* |
| *D7 to 14* | *0.01* | *0.01* | *0.48* | *0.00* |
| *D14 to 28* | *0.01* | *0.01* | *0.71* | *0.00* |
| *D28 to 35* | **0.06** | **0.01** | **< 0.00** | **0.00** |
| *D35 to 42* | *0.12* | *0.02* | *0.48* | *0.00* |
| *D42 to 49* | *0.02* | *0.02* | *0.52* | *0.00* |
| *D0 to 49* | *0.07* | *0.07* | *0.26* | *0.00* |
| **FCR (kg/pen)** |  |  | **Mean** | **SEM** |
| *Control (n=17)* |  |  | 2.34 | 0.03 |
| *+PE fiber (n=20)* |  |  | 2 | 0.14 |

We also observed significant differences in BWG for the ternary mixture treatment groups from D7 to D14, D14 to D28, D28 to D35 and D42 to D49 (Table 3). Analysis of the mean FCR of control, low dose and high dose mixture treatment groups over the treatment period (49 days) indicated that low dose mixture and high dose mixture treatment groups have a lower mean FCR compared to the control group (Table 3). FI was also assessed to evaluate feed consumption over the treatment period. FI increased significantly over the treatment period for control, low dose and high dose groups (Figure S2). However, at D28 for low dose and high dose treatment groups is significantly increased compared to the control group (Figure S2, P < 0.001).

**Table 3.** Summary of body weight gain and feed conversion ratio results comparing control, low dose mixture and high dose mixture treatment groups. ^a^.

|  |  |  |  |  |  |  |
| --- | --- | --- | --- | --- | --- | --- |
| **BWG (kg/bird)** | **Control mean (n=17)** | **Low dose mixture mean (n=22)** | **High dose mixture mean (n=19)** | **P-value <sup>a</sup>** | **F\*ratio <sup>b</sup>** | **dFn, dFd <sup>c</sup>** |
| D0 to 7 | 0.004 | 0.003 | 0.003 | 0.87 | 0.14 | 2, 51.99 |
| D7 to 14 | 0.009 | 0.007 | 0.012 | **0.00** | 7.39 | 2, 53.90 |
| D14 to 28 | 0.004 | 0.013 | 0.013 | **< 0.00** | 162 | 2, 45.22 |
| D28 to 35 | 0.069 | 0.011 | 0.011 | **< 0.00** | 313 | 2, 22.38 |
| D35 to 42 | 0.019 | 0.018 | 0.017 | 0.42 | 0.87 | 2, 52.53 |
| D42 to 49 | 0.017 | 0.021 | 0.018 | **0.02** | 4.15 | 2, 50.32 |
| D0 to 49 | 0.067 | 0.071 | 0.018 | **< 0.00** | 365 | 2, 44.62 |
| **FCR (kg/pen)** |  |  |  |  | **Mean** | **SEM** |
| Control (n =17) |  |  |  |  | 2.34 | 0.03 |
| Low dose mixture (n=22) |  |  |  |  | 1.87 | 0.26 |
| High dose mixture (n=19) |  |  |  |  | 1.90 | 0.24 |
| <sup>a</sup>. Bolded values indicate significance (Brown-Forsythe ANOVA test; P-value ≤ 0.05).<br><sup>b</sup>. The P-value is computed from the F* ratio and is analogous to the F ratio commonly used for ANOVA.<br><sup>c</sup>. Degrees of freedom for the numerator (dFn) and denominator (Dfd) used for calculating the F*ratio. |  |  |  |  |  |  |

Histological evaluation was then conducted to evaluate if pathological damage occurred following exposure to the +PE fiber, low dose, and high dose treatment groups. There were no overt signs of inflammation, lesions, abnormal cellular development, or other significant changes to the intestinal tract in any group (Figure S3- S4). In addition, mortality rates over the trial did not appear to be correlated to treatment due to several of the control group mortalities being attributed to wing band injuries early in the study period and heat stress at later stages of the trial. However, the high dose group did have four mortalities reported from D7-D14 unlike the low dose group which had zero from D7-D42 (Figure S5).

### 3.2. Cecal microbiome significantly altered by PE fiber MPs presence

To assess the effect of PE fiber MPs on the cecal microbiome composition, the control (untreated) group was compared to the PE fiber MPs group. To assess this, 16S rRNA gene amplicon sequencing targeting the V4 region was conducted. The top 50 microbial taxa were plotted to the genera level (Figure 1). Overall, there were differences between samples across treatment groups (Figure 1). The main effect “treatment” was not significant for α-diversity metrics, indicating that phylogenetic diversity within samples was not statistically different (Table S3) (Kruskal-Wallis pairwise, P-value ≥ 0.05 Q-value ≥ 0.05), However, overall β-diversity metrics comparing control to +PE Fiber were statistically significant as determined by PERMANOVA (Table S4) (P-value ≤ 0.05;Q-value ≤ 0.05). Thus, confirming phylogenetic differences between the control and +PE Fiber treatment groups. Overall assessment of dissimilarities among the treatment groups as determined by Bray-Curtis and Weighted UniFrac PCoA plots also demonstrated distinct clustering (Figure S6).

**Figure 1.**
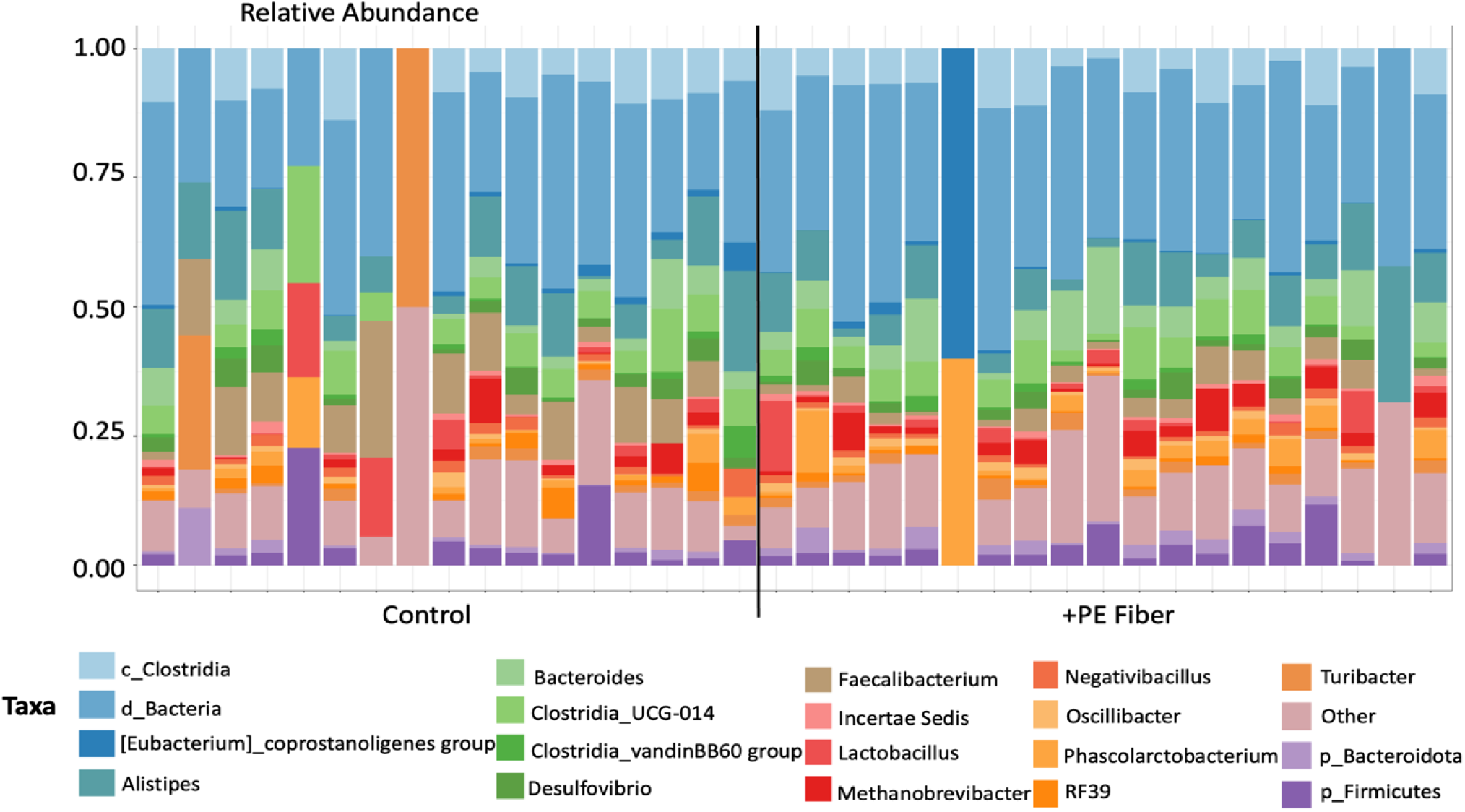
The relative abundance of the top 50 genera in each sample separated by treatment group (black line).

Differential abundance testing using ANCOM-BC was subsequently conducted to quantify changes in relative abundance of microbial taxa as related to the control group. It was determined that genera, *Fournierella,* and an unclassified genus in the family, *Coriobacteriaceae,* were enriched in the +PE Fiber treatment group (Figure 2). While the genus *Synergistes,* and one unclassified genus in the *Desulfovibrionaceae* family, were depleted in +PE Fiber cecal microbiomes (Figure 2). We subsequently assessed microbial composition variability among control and +PE fiber treatment groups at the phylum level. Analysis of the mean relative abundance of phylum was used to evaluate potential dysbiosis in the cecal microbiomes. The mean relative abundance of *Firmicutes* was higher in the control group while *Bacteroidota* was higher in the +PE Fiber treatment group, indicating (Figure 3). Indicating the +PE Fiber treatment group had a lower *Firmicutes/Bacteroidota* ratio compared to the control treatment group (49). These results also highlighted that the +PE Fiber treatment group had a higher mean relative abundance of *Bacteroidota*, and *Desulfobacterota,* while the control group exhibited a higher mean relative abundance of *Firmicutes* and *Proteobacteria* (Figure 3).

**Figure 2.**
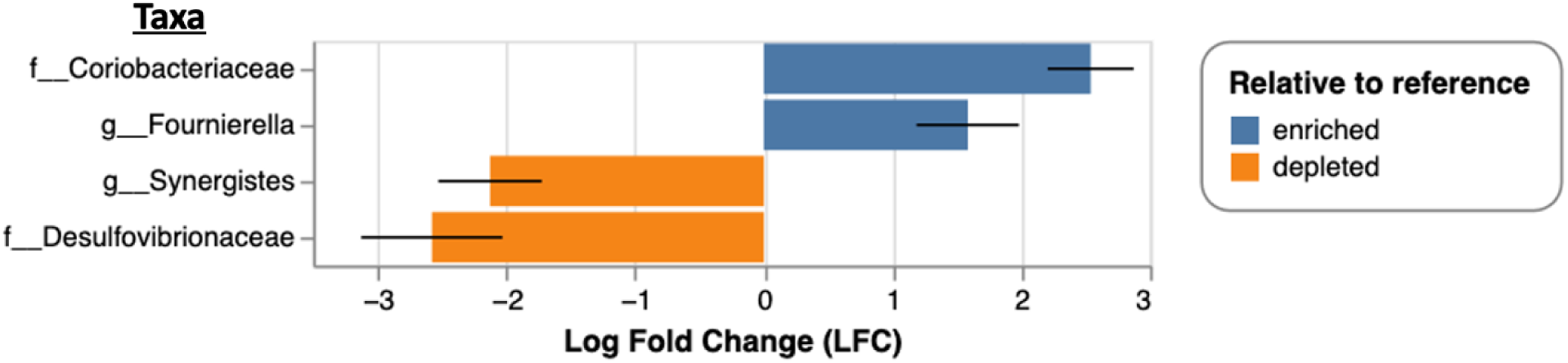
Differential abundance of genera in the +PE Fiber group relative to the control group as determined by ANCOM-BC. Unclassified genera were annotated to the family level.

**Figure 3.**
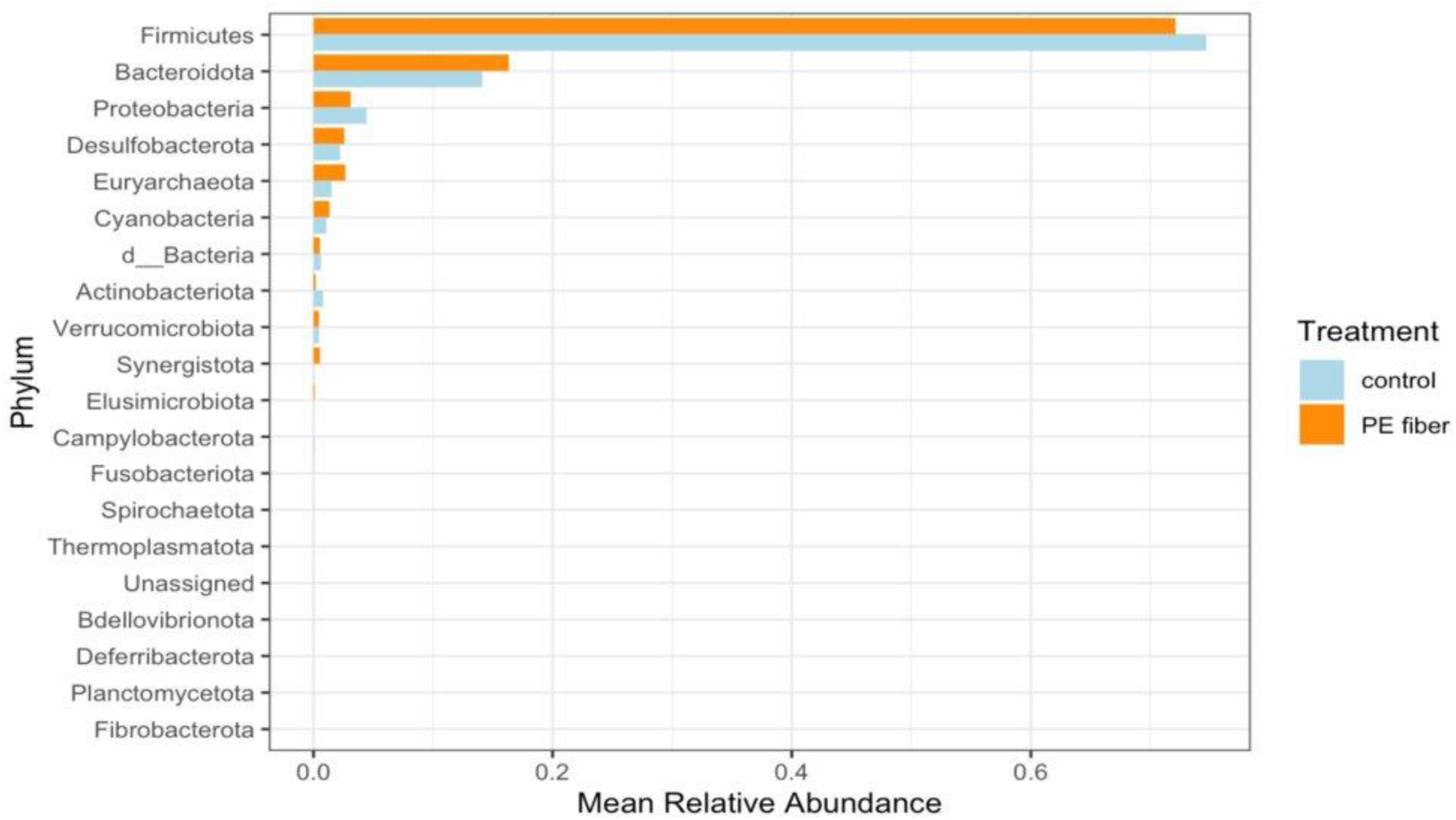
Comparison of mean relative abundance of phylum by treatment group for +PE fiber trial.

### 3.3. Ternary mixtures of agricultural chemicals significantly alter cecal microbiomes

In this study, 16s rRNA amplicon sequencing was utilized to evaluate changes in the cecal microbial composition following exposure to low and high concentrations of a ternary mixture of nitrate, atrazine and imidacloprid. Overall, the cecal microbiomes of each treatment group exhibited an abundance in *Firmicutes* and *Bacteroidota* (49).(Figure 4). This corresponds with our previously sequenced poultry microbiomes (24, 50, 51). In addition, broilers in the low dose mixture and high dose mixture treatment groups had a higher relative frequency of *Euryarchaeota* (49).(figure 4).

**Figure 4.**
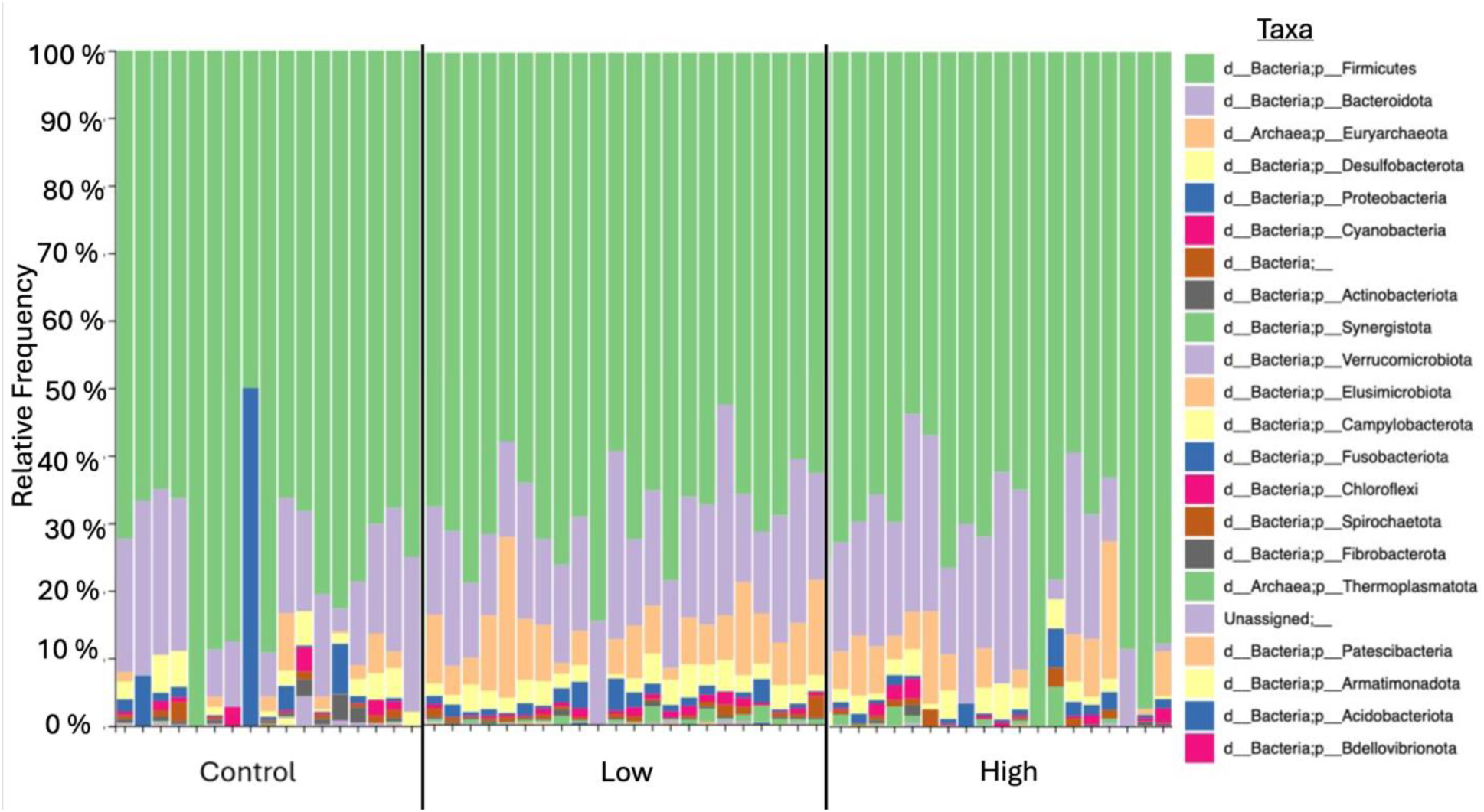
Taxa bar plot showing the relative frequency of phyla by treatment group. Treatment groups are separated by a solid black line.

To assess treatment effects on microbial composition, ANOVA for α-diversity metrics were evaluated. The results indicated that the main effect “treatment” (low dose and high dose ternary mixtures) significantly impacted microbial diversity and richness (Table S5) (P-value ≤ 0.05). Further evaluation of treatment effects on microbial composition was conducted using Kruskal-Wallis pairwise comparisons. Comparison of the control and high dose mixture treatment groups detected significance for α-diversity metrics, Faith’s phylogenetic diversity and observed features (Table S5) (P-value ≤ 0.05; Q-value ≤ 0.05). However, Kruskal-Wallis pairwise results for α-diversity metrics were not significant for comparisons of the control group and low dose mixture treatment groups (Table S6) (P-value ≥ 0.05; Q-value ≥0.05). Pairwise comparison of the high dose mixture treatment group to the low dose mixture treatment group did yield significance for Shannon’s diversity index, Faith’s phylogenetic diversity and observed features (Tables S6) (P-value ≤ 0.05; Q-value ≤ 0.05).

We then evaluated treatment effects as related to β-diversity metrics. First, PERMANOVA was used for β-diversity metrics to assess significant differences in taxonomic composition between groups. The main effect “treatment” was determined to significantly impact cecal microbial composition based on β-diversity metrics (Table S7) (P-value ≤ 0.05; Q-value ≤ 0.05). Significance was detected for all comparisons (Tables S7). The results were subsequently visualized using principal coordinate analysis in Emperor. PCoA plots provided a visual indication of similarities and dissimilarities among the treatment groups. It was determined that the low and high dose mixture treatment groups clustered more closely together indicating similarity in microbial composition (Figure S6).

Next, differentially abundant taxa were determined using ANCOM-BC. Differential abundance analysis as related to the control group revealed that the genera *Akkemansia*, *Fournierella*, *Ruminococcus* and an unclassified genus in the family *Coriobacteriaceae* were enriched in the low dose and high dose mixture treatment groups (Figure 5). One taxon was depleted in the control group which was identified as an unclassified genus in the family *Desulfovibrionaceae.* This data further confirmed the dissimilarity between the control and ternary mixture treatment groups.

**Figure 5.**
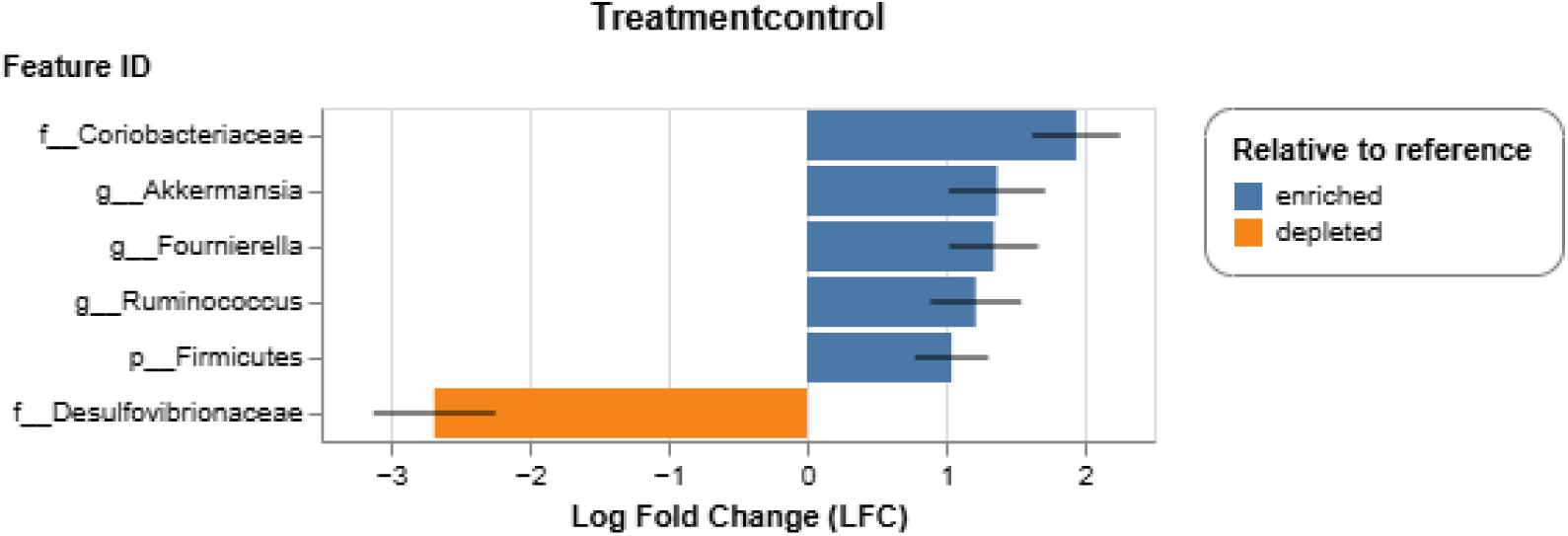
Differential abundance of genera in the low dose and high dose groups relative to the control group as determined by ANCOM-BC. Unclassified genera were annotated to the family level.

### 3.4. PE fiber MPs significantly altered cecal metabolomes of broilers

To assess the impact PE fiber MPs have on the cecal metabolome, a pairwise comparison was conducted on the total metabolome of each group (i.e, control cecal metabolome vs +PE Fiber cecal metabolome). Analysis of the total metabolome as determined with an unsupervised model, principal component analysis, and a supervised model, partial least square discriminant analysis (PLS-DA), highlighted distinct clustering of the total metabolome of +PE fiber (Figure 6). A summary of all detected metabolites is provided in Table S8. Results of pairwise comparisons indicated there were 113 metabolites that were significantly down-dysregulated, 73 significantly up-dysregulated and 4,742 insignificant metabolites (Table S9). Volcano plot analysis was then conducted in Compound Discoverer which highlighted that Vitamin C, Glucuronolactone, 2-arachidonoylglycerol, and dihydroxyphenylalanine were among the metabolites significantly down dysregulated in the +PE Fiber treatment group (Table S10) (Log2 fold change ≤ -2; P-value ≤ 0.05). While associated metabolites for the control treatment group included Uplandicine, Ethyl docosahexaenoate, and methyl isoquinoline-3-carboxylate as annotated by Compound Discoverer (Table S10) (Log2 fold change ≥ 2; P-value ≤ 0.05). Overall, there were a greater number of metabolites dysregulated due to the presence of +PE Fiber compared to the control. These results indicate that the presence of PE fiber significantly impacts cecal microbial activity.

**Figure 6.**
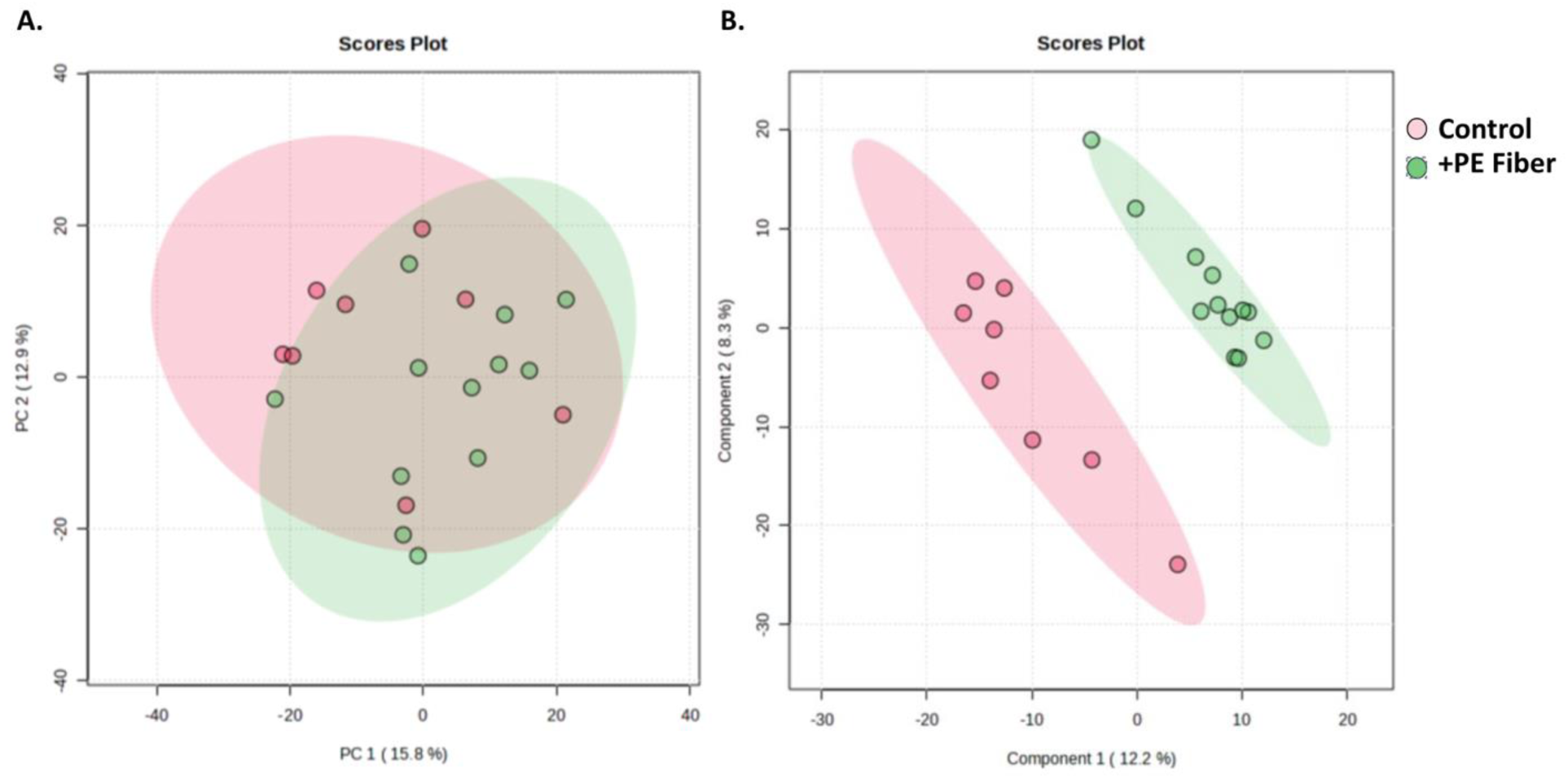
Analysis of the total cecal metabolomes of control, and +PE Fiber treatment groups with (A) an unsupervised principal component analysis model and (B) partial least square discriminant analysis (PLS- DA).

Next, pathway-level changes in the +PE fiber treatment group were assessed using the functional analysis module in MetaboAnalyst. It was determined that there was significant modulation in metabolic pathways associated with the +PE fiber treatment group. Specifically, there was a higher total ion intensity of metabolites linked to the pentose phosphate pathway, pentose and glucuronate interconversion, primary bile acid biosynthesis, steroid biosynthesis, and sphingolipid metabolism (Figure 7). In the +PE Fiber metabolome there was also a noticeably higher intensity of features 175.152_81.21 *m/z*_RT, 174.1486 102.92 *m/z*_RT, 192.1592_84.47 *m/z*_RT, 193.1625_84.47 *m/z*_RT (Figure 7). For each feature the *m/z* represented mass to charge ratio, and RT is retention time in seconds for that metabolite feature. These metabolites could not be mapped to metabolic pathways. Despite this, the dysregulation of these metabolites further highlights that PE fiber MPs significantly impacted microbial activity.

**Figure 7.**
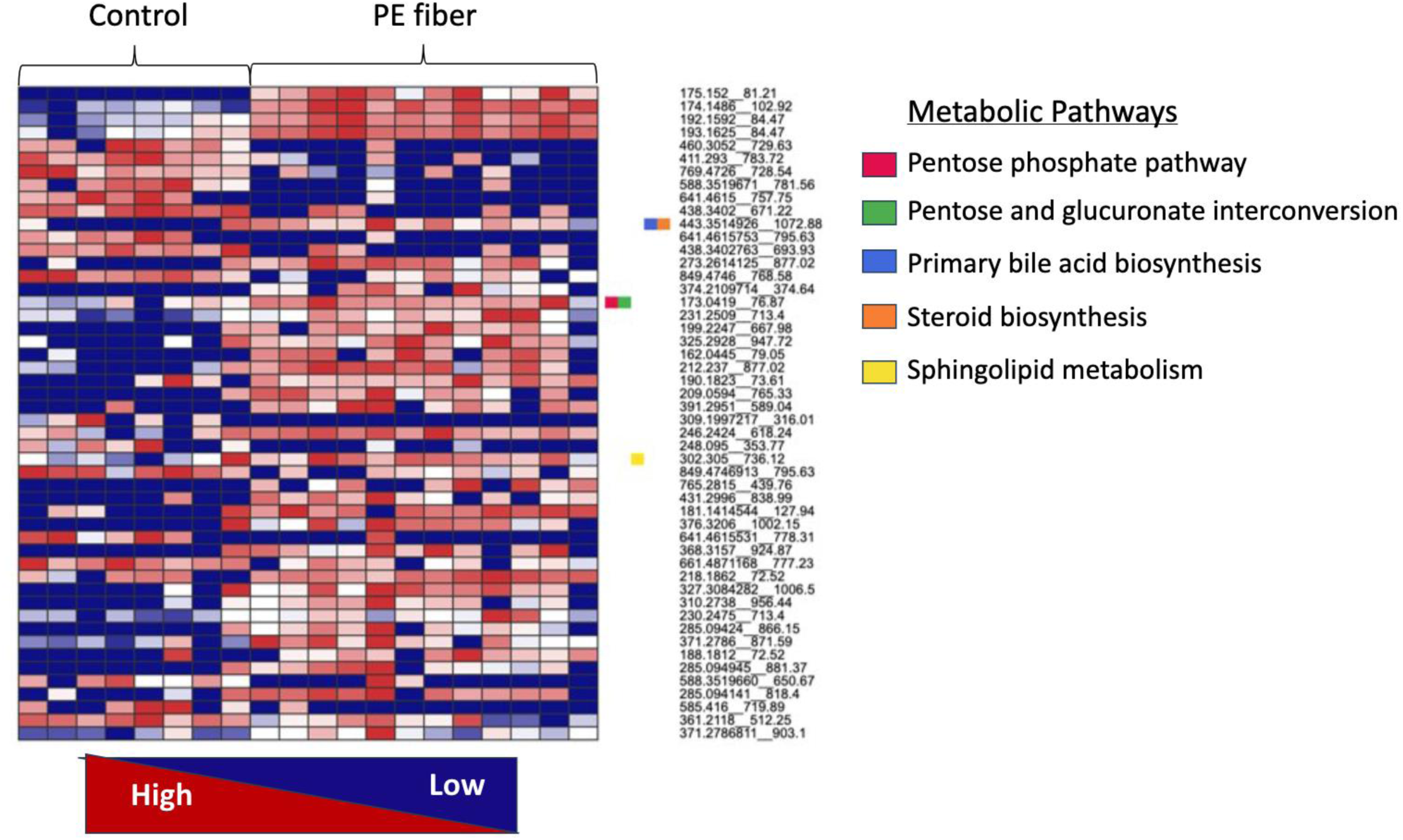
Pathway-level analysis performed in MetaboAnalyst 6.0 using the Function analysis module. Functional analysis incorporates a Gene Set Enrichment and Mummichog algorithm (P-value = 0.05, KEGG library *Gallus gallus* library) for accurate detection of changes in total ion intensity of features.

### 3.5. Cecal metabolome significantly altered by ternary mixtures in a dose-dependent manner

To determine metabolome similarities and dissimilarities between control and treated metabolomes (low dose and high dose ternary mixtures), a multi-group comparison of the total metabolomes was conducted. Multiple group analysis of the total metabolome as determined with principal component analysis highlighted distinct separation of the total metabolomes of control group from the ternary mixtures (Figure 8). Analysis of variance analysis (ANOVA) was subsequently conducted to investigate differences in metabolites among the treatment groups. ANOVA analysis detected twelve significant metabolites of which Methylisopelletierine and Procarbazine were putatively identified (Table 4). Neither metabolite was detected in the annotated features list derived from Compound Discoverer (Table S12).

**Figure 8.**
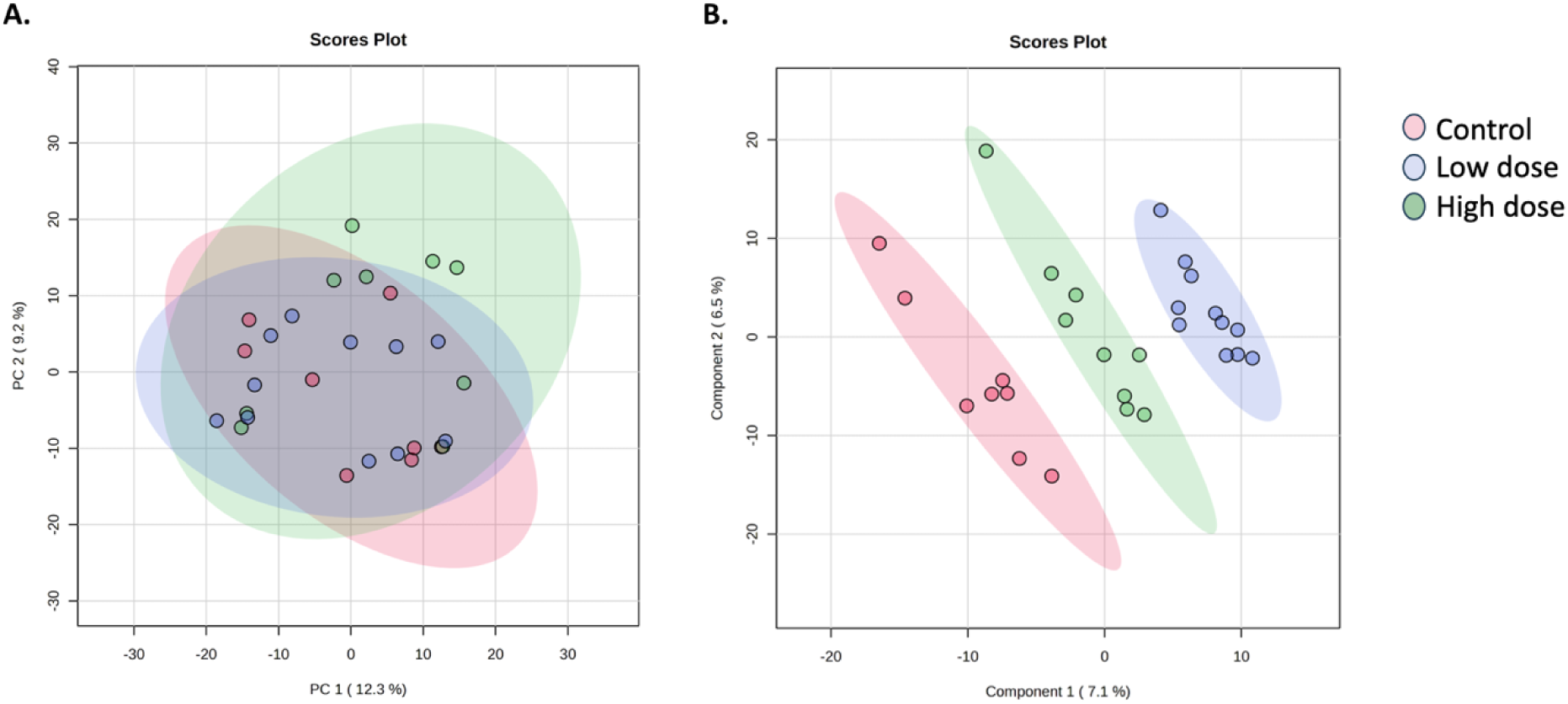
Analysis of the total cecal metabolomes of control, low dose and high dose treatment groups with (A) an unsupervised principal component analysis model and (B) partial least square discriminant analysis (PLS-DA)).

**Table 4.** Significant metabolites as determined using ANOVA (Tukey’s post-hoc test; P-value ≤ 0.05)

|  |  |  |  |  |  |
| --- | --- | --- | --- | --- | --- |
| *m/z*_RT(s) | *Putative ID* | *F-value* | *P-value* | *-Log10(p)* | *FDR* |
| 175.1519__86 | Not available | 54.133 | 8.2266e-10 | 9.0848 | 1.6432e-06 |
| 174.1486__103.39 | Not available | 52.677 | 1.0845e-09 | 8.9648 | 1.6432e-06 |
| 192.1592__82.74 | Not available | 52.038 | 1.2265e-09 | 8.9113 | 1.6432e-06 |
| 581.7156__145.79 | Not available | 51.607 | 1.3337e-09 | 8.8749 | 1.6432e-06 |
| 174.1486__83.83 | Not available | 44.409 | 5.911e-09 | 8.2283 | 5.8259e-06 |
| 193.1625__82.74 | Not available | 43.146 | 7.8245e-09 | 8.1065 | 6.4265e-06 |
| 156.1381__82.74 | Methylisopelletierine | 36.603 | 3.7373e-08 | 7.4274 | 2.6311e-05 |
| 568.2721__125.15 | Not available | 28.857 | 3.1957e-07 | 6.4954 | 0.000197 |
| 244.1406__79.47 | Procarbazine | 25.552 | 9.0512e-07 | 6.0433 | 0.000496 |
| 555.2878__273.39 | Not available | 18.71 | 1.0781e-05 | 4.9674 | 0.005313 |
| 242.1464__611.08 | Not available | 15.646 | 3.9243e-05 | 4.4062 | 0.017581 |
| 366.9677__71.82 | Not available | 13.474 | 0.000107 | 3.9704 | 0.043965 |

Hierarchical clustering was used to evaluate the cecal metabolomes for similarities and dissimilarities of detected dysregulated metabolites within each treatment group. We observed dose-response changes to the cecal metabolomes exposed to both ternary mixtures (Figure 9). Putatively identified metabolites as determined by hierarchical clustering included Heptyl 4-hydroxybenzoate, Procarbazine, Ganglioside GD3 d18, and Methylisopelletierine (Figure 9). Heptyl 4-hydroxybenzoate had a higher intensity in the control and high dose mixture treatment groups (Figure 9). Procarbazine and Ganglioside GD3 d18 both had a lower intensity in control and high dose cecal metabolomes (Figure 9). Both the low dose and high dose cecal metabolomes had a higher intensity of Methylisopelletierine (Figure 9). A dose response was also detected for features, 230-1172_478.09, 370.1431_473.76, and 242.1464_478.611.08 (Figure 9). Due to the mass spectrometry methods selected nitrate was not able to be detected (Tables S12-13). However, atrazine but not imidacloprid was detected in the annotated peak list in Compound Discoverer suggesting more rapid metabolism of atrazine (Tables S12). In all, these results further highlight the significant difference in total ion intensity of individual metabolites between the treatment groups.

**Figure 9.**
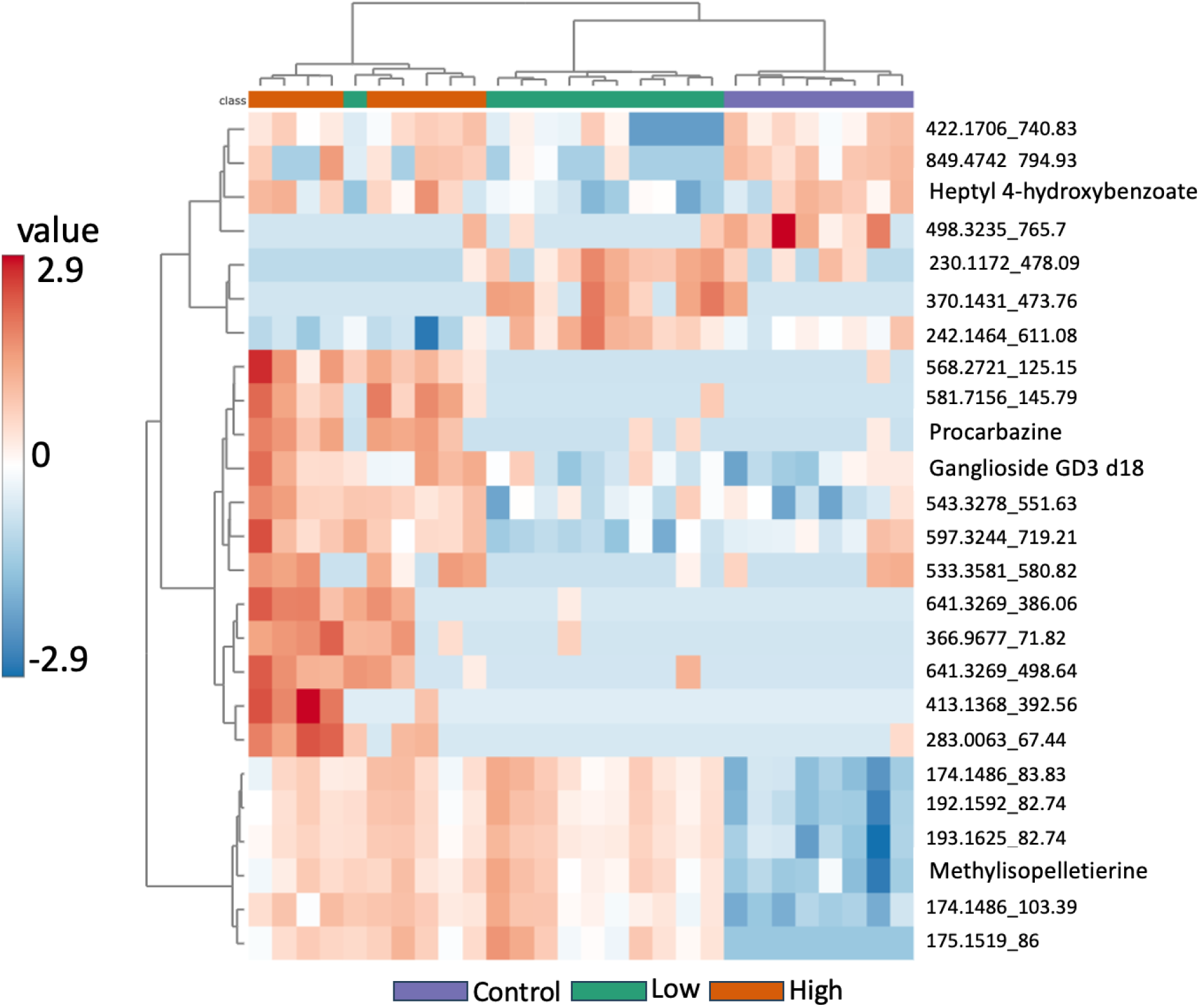
Hierarchical clustering analysis results of the top 25 metabolites detected among the treatment groups. Colored boxes (red to blue) represent the normalized total ion intensity of metabolites within samples.

Functional analysis (P-value = 0.05, KEGG *Gallus gallus* library) was then conducted to assess pathway-level changes to the cecal metabolomes. Within MetaboAnalyst significant hits refers to the number of empirical compounds in a data set detected in a specific metabolic pathway. The following metabolic pathways contained significant feature hits: metabolism of xenobiotics by cytochrome P450, amino sugar and nucleotide sugar metabolism, pyruvate metabolism, thiamine metabolism and pantothenate and CoA biosynthesis (Table 5). Additionally, metabolika pathway analysis in Compound Discoverer determined key pathways within the cecal metabolomes such as the superpathway of glycolysis, pyruvate dehydrogenase, tricarboxylic acid (TCA), and glyoxylate bypass, and the superpathway of acetyl-CoA biosynthesis (Table S13). Given the results from the untargeted metabolomics statistical analyses, it can be concluded that the poultry cecal metabolome is significantly modulated following exposure to both ternary mixtures.

**Table 5.** Detected metabolic pathways (P-value = 0.05, KEGG *Gallus gallus* library)

| Metabolic Pathway | Hits - total | Hits-significant | AdjP - Fisher | AdjP - EASE | AdjP - Gamma | Compound Hits |
| --- | --- | --- | --- | --- | --- | --- |
| Metabolism of xenobiotics by cytochrome P450 | 26 | 2 | 0.24 | 1 | 0.06 | C14851;C14850;C14556;C14874;C14849 |
| Amino sugar and nucleotide sugar metabolism | 30 | 1 | 0.39 | 1 | 1 | C03410 |
| Pyruvate metabolism | 4 | 1 | 0.24 | 1 | 1 | C03451 |
| Thiamine metabolism | 1 | 1 | 0.08 | 1 | 1 | C00378 |
| *Pantothenate and CoA biosynthesis* | 8 | 1 | 0.24 | 1 | 1 | C00831 |

## 4. DISCUSSION

In this study we first aimed to elucidate the effects of PE fiber MPs on the cecal microbiome and metabolome of broiler chickens. The data presented demonstrates that the presence of PE fiber MPs does significantly alter cecal taxonomic composition and cecal microbial activity. It has been well established that MPs result in adverse health outcomes in various species (52–55). However, there is a lack of research on the impact this emerging contaminant has on terrestrial food protein sources such as poultry. Our previous study reported that PE fiber exposure resulted in less *Firmicutes* than *Bacteroidetes* in an *in vitro* cecal mesocosm (24). A similar trend is also observed in this exposure study. We also observed with differential abundance analysis that *Fournierella* was enriched in the +PE Fiber treatment group while *Synergistes* were depleted. A similar increase in *Fournierella* was observed in an *in vitro* cecal PolyFermS model (21). In that study, Asare et al. (2021) (21) observed that Viande Levure medium supplemented with 2.5 g/L of fructooligosaccharides (FOS), a common prebiotic (56, 57), increased the abundance of *Fournierella* and *Pseudomonas.* In contrast, Jin et al. (2024) (58) observed an increased abundance of *Fournierella* correlated to reduced intestinal calcium. In this study, the +PE Fiber group also presented with significantly lower BWG and FI on D28 compared to control birds (Table 2) further indicating the possibility of poor nutrient absorption, loss of appetite or gut dysbiosis. Given this, the presence of PE fiber may elicit similar microbial composition changes as FOS and subsequently inhibit intestinal absorption in poultry.

We also detected the depletion of *Synergistes* in the +PE Fiber treatment group. Wang et al. (2023) (59) noted that cecal microorganisms *Synergistes* and *norank_f_Desulfovibrionaceae* were correlated with high levels of inflammatory cytokine, IL-6, which negatively impacted growth. It has also been seen that probiotics can alter cytokine expression (60, 61). Nonetheless, the perturbation to cecal microbial composition occurred despite having a slight BWG difference among the control and +PE fiber groups aside from D28. Considering the depletion of this genera and growth parameters for the +PE Fiber group, it is plausible that PE fiber presence did not induce an inflammatory response. However, future studies should explore if PE Fiber presence impacts cytokine expression in broilers via modulation of cecal microbiome composition.

Pathway-level analysis highlighted that the presence of PE fiber MPs significantly impacted features linked to the pentose phosphate pathway, pentose and glucuronate interconversion, primary bile acid biosynthesis, steroid biosynthesis, and sphingolipid metabolism. Analogous to this study, Teng et al. (2021) (62) demonstrated that an ecologically relevant concentration of PE (10 μg/L) induced modulation of sphingolipid metabolism and the pentose phosphate pathway in oysters. Sphingolipids are a class of lipid molecules that are essential in regulating eukaryotic membranes and cell function (63, 64). Within sphingolipid metabolism, ceramides are routinely produced via sphingolipid degradation (64). The increased production of ceramides has been correlated with various metabolic diseases (64). Considering this, the dysregulation of sphingolipid metabolism in the +PE Fiber treatment group indicates these broilers may be at an increased risk of developing metabolic diseases.

Previous research by Wen et al (2024) (65) demonstrated that oral exposure to environmentally relevant concentrations of polystyrene MPs in mice induced intestinal damage and dysregulated bile acid profile in feces. In this study, we also noted significant dysregulation of bile acid metabolism but in the cecal metabolome following PE MP exposure. Bile acids are cholesterol derivatives produced in the liver that act as amphipathic surfactants and regulate macronutrients (i.e., lipids, carbohydrates and proteins (66, 67). Bile acids are also essential for maintaining glucose and lipid homeostasis (67). Disruption of the homeostasis of bile acids has been shown to negatively impact health by inducing metabolic disorders (68, 69). In all, the data presented here may indicate that PE fiber MPs are inducing bile acid metabolism disorders in broilers.

Next, we investigated the impact known groundwater contaminants, agricultural chemicals, have on poultry microbiomes and metabolomes. Agricultural chemicals are continuously contaminating waterways typically via atmospheric deposition, and agricultural runoff or erosion events (70, 71). Given that agricultural chemicals are being detected above safe standards (23), chemical risk assessment must be modified to account for these complex mixtures. Here, we aimed to elucidate the impacts of an ecologically relevant mixture of nitrate, atrazine and imidacloprid using a low dose mixture and a high dose mixture. We selected a mixture of nitrate, atrazine and imidacloprid based on their frequency of detection and concentrations exceeding enforcement standards in the 2021 DATCP Targeted Sampling Report (23). Our previous study indicated that exposure to the low dose mixture and two other ternary mixtures greatly altered the growth kinetics of poultry cecal microbiomes indicating potential chemical-biological effects from this ternary mixture (29). We also observed that a two-chemical combination of these chemicals at 3,000 ppb elicited a greater decline in Caco-2 cell viability (29). Biological impacts were also detected in this *in vivo* study with poultry cecal microbiome samples exposed to the same low dose ternary mixture as evidenced by varied cecal microbial composition in this treatment group (Figure 1). We further demonstrated that the low dose mixture (35,000 ppb nitrate + 1.7 ppb atrazine + 0.58 ppb imidacloprid) and the high dose mixture (100,000 ppb nitrate + 3,000 ppb atrazine + 3,000 ppb imidacloprid) exposure resulted in increased abundance on five taxa not enriched in the control group (Figure 5). The ternary mixtures used in this study also significantly altered microbial activity as determined using hierarchical clustering analysis of the metabolomics results (Figure 5).

Overall, we did not observe overt signs of toxicity or treatment related mortalities to the broilers, although significant differences in BWG and FI were observed. Foster & Khan (1976) (72) reported that laying hens exposed to 100 ppm atrazine also did not exhibit any physiological changes or signs of toxicity, but that atrazine residues persisted in the fecal excrement for up to 4 days after the exposure period ended. They concluded that the continuous presence of atrazine and related degradation products were evidence of bioaccumulation in tissues (72). Therefore, bioaccumulation in tissues, especially if occurring in the gastrointestinal tract, may explain the modulations to the cecal microbial composition observed in this study. Specifically, we observed the enrichment of *Akkermansia*, *Fournierella* and *Ruminococcus* in the treated groups. It cannot be determined which taxa were altered by either nitrate, atrazine or imidacloprid. However, correlations can be inferred based on current literature. For example, an *in vivo* study demonstrated that frogs exposed to atrazine experienced perturbations to metabolic pathways pantothenate and CoA biosynthesis, pyruvate metabolism and pyruvate metabolism (73). Likewise, we noted these three pathways had significant feature hits as determined with MetaboAnalyst 6.0.

In addition, we observed that thiamine metabolism had a significant feature hit. Both pantothenate and thiamine are B vitamins which act as cofactors for enzymes to regulate cellular and mitochondrial energy metabolism (74). Thiamine specifically is involved in the activation and maintenance of immune cells and protein (75). In humans, thiamine deficiency results in impairments to the nervous system, energy metabolism, and digestion among other essential metabolic functions (75). Poultry do not naturally produce thiamine so a diet deficient in this B vitamin can result in low carcass weight, low hatchability, compromised carbohydrate, protein, and lipid metabolism or mortality (76). Thiamine is supplemented in the poultry diet used in this study, so its depletion is indicative of consumption or utilization in the gastrointestinal tract of the broilers. Thus, emphasizing the risks of modulation to pantothenate, and CoA biosynthesis and thiamine metabolism pathways as seen in this study.

The results presented in this study highlight the impact of a long-term exposure (49 days) of environmental contaminants to broiler health which may have potential implications on human populations. It is well-known that pesticide and MP exposures pose significant human health risks (3, 5). However, due to differences in species sensitivity, exposure duration (e.g., acute versus chronic), body mass and several other factors, it becomes complex to understand the full extent of said risk. For instance, Witwicka et al. (2025) (77) demonstrated in bumble bees that acute or short-term exposure (12 days) to neonicotinoid pesticides upregulates more apoptosis-inducing genes compared to chronic or long-term exposure (12 days) which suggests that acute exposure induces immediate lethal impacts. Whereas both acute and chronic atrazine exposure in female zebrafish has been documented to alter reproductive development (78–81) and impair the endocrine system (82, 83). In this study, it was also evident that chemical concentration had varied impacts on the poultry cecal microbiome and metabolome. However, it is unclear if or how biotransformation of nitrate, atrazine and imidacloprid impacts broiler health given that atrazine appears to bioaccumulate more easily in poultry. Future studies should also elucidate the impacts of these contaminants on inflammatory cytokine expression as related to the cecal microbiome which may allow for targeted approaches to reduce incidences of gut dysbiosis or poor growth parameters.

## 5. CONCLUSIONS

In this study, we provide evidence of PE fiber MPs modulating cecal microbiome composition and microbial activity following a long-term exposure. 16S rRNA amplicon sequencing indicated that β-diversity metrics comparing control to +PE Fiber treatment group were statistically significant. Interestingly, the +PE Fiber group was enriched in *Fournierella,* but depleted in *Synergistes* indicating potential beneficial impacts to PE fiber presence. However, untargeted metabolomics results highlighted significant dysregulation of metabolic pathways in the +PE Fiber treatment group which are associated with increased incidence of metabolic disorders. Further, we demonstrate how a ternary mixture of nitrate, atrazine and imidacloprid, even at environmentally relevant concentrations, are capable of modulating poultry cecal microbial composition and cecal microbial activity. Thus, highlighting that agricultural chemicals and the emerging contaminant, MPs, impact health outcomes to broilers and may subsequently impact consumers’ health. These results underscore the importance of evaluating agricultural chemicals as mixtures and at environmentally relevant concentrations.

Overall, the data presented here provides evidence of the potential health risks to livestock and humans following exposure to environmentally relevant concentrations of PE fiber MPs and agricultural chemical mixtures.

## Acknowledgements

The authors would like to thank the University of Wisconsin-Madison’s Poultry Research Facility and staff members for managing the husbandry of the broilers. We would also like to thank Dr. Steve Ricke and the following members of the Ricke laboratory who provided support during this study: Ashley A. Tarcin, Jessica Brown, Margaret Costello, Colin Wallirich, Maxim A. Peckenschneider, Haley Tarcin, and Emily Matiak. The authors also thank Dr. Bradley Bolling and Klay Liu for providing access to the mass spectrometer used for untargeted metabolomics analysis. In addition, the author C.C. would like to acknowledge Fuad Shatara for preparing the polyethylene fiber microplastics utilized in this study and Gordon Winkler for their support during this study.

## Data Availability

Sequencing data were uploaded to NCBI’s Sequence Read Archive as BioProject ID: PRJNA1200004 (https://www.ncbi.nlm.nih.gov/bioproject/PRJNA1200004). Untargeted Metabolomics data was uploaded to MetaboLights as project MTBLS11958 (https://www.ebi.ac.uk/metabolights/MTBLS11958).

## Supplemental Figures and Tables

**Figure S1.**
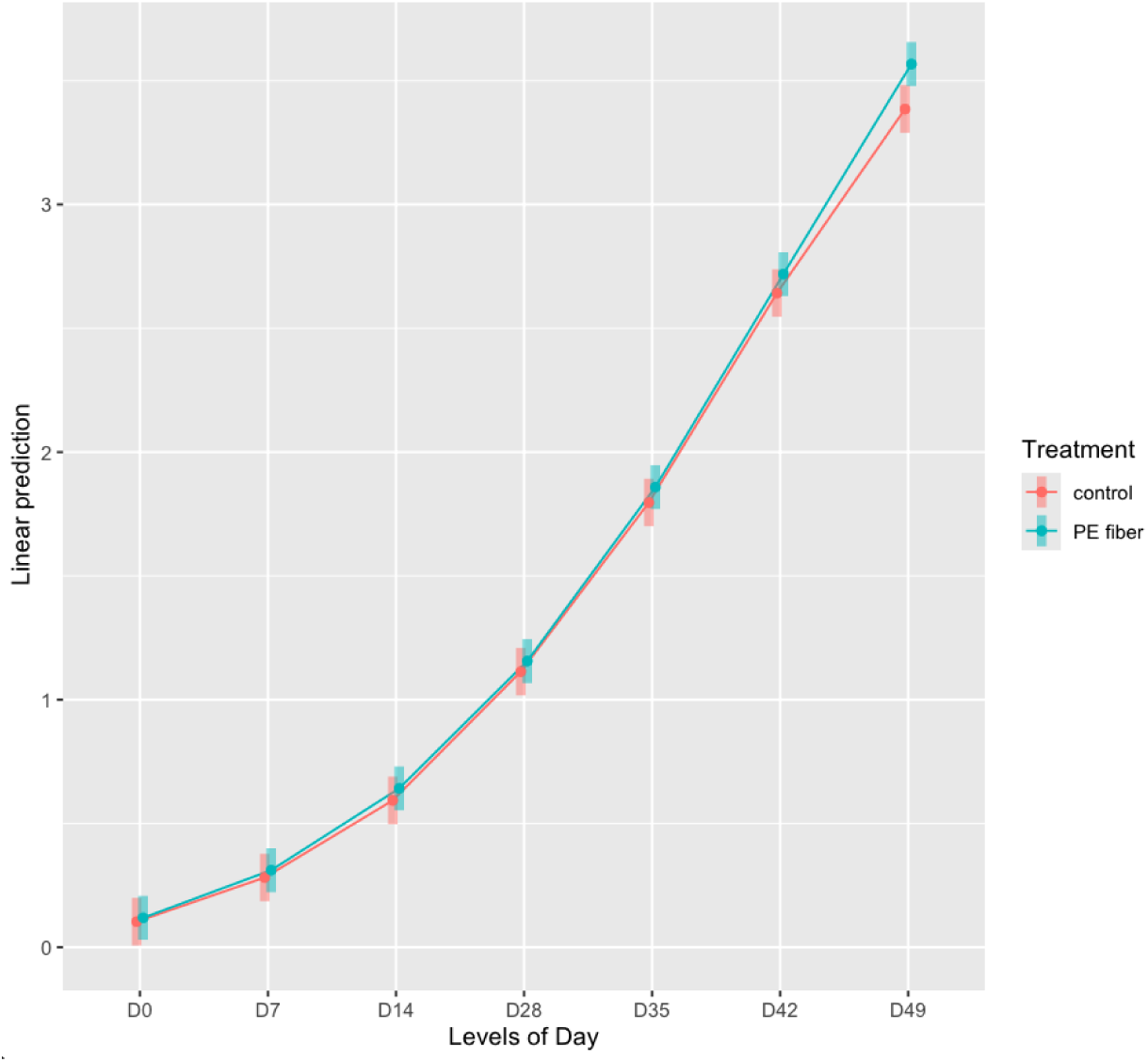
Linear Mixed Effect Model analysis assessing feed intake for control and +PE Fiber treatment groups (P ≤ 0.05).

**Figure S2.**
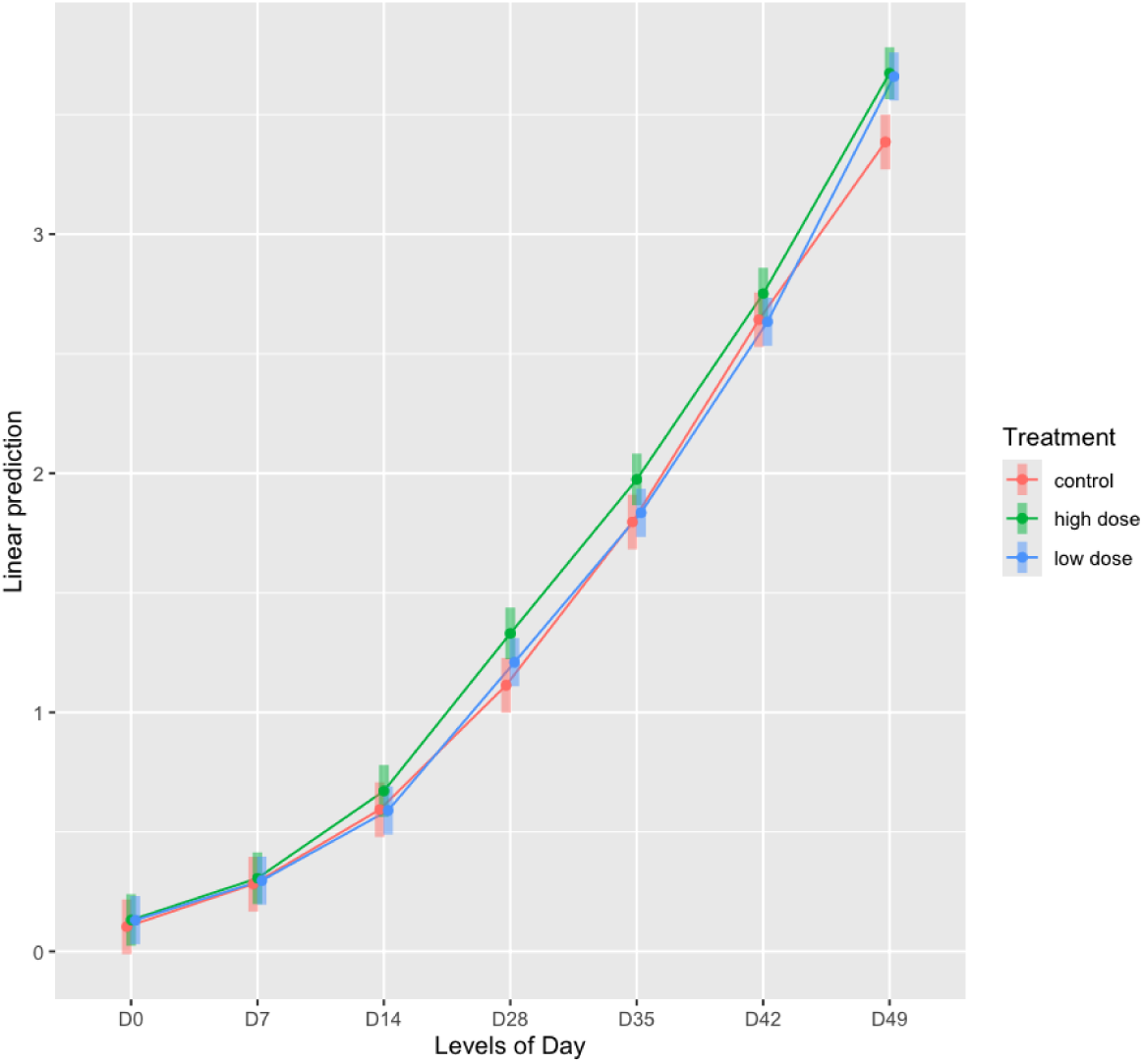
Linear Mixed Effect Model analysis assessing feed intake for control, low dose, and high dose treatment groups (P ≤ 0.05).

**Figure S3.**
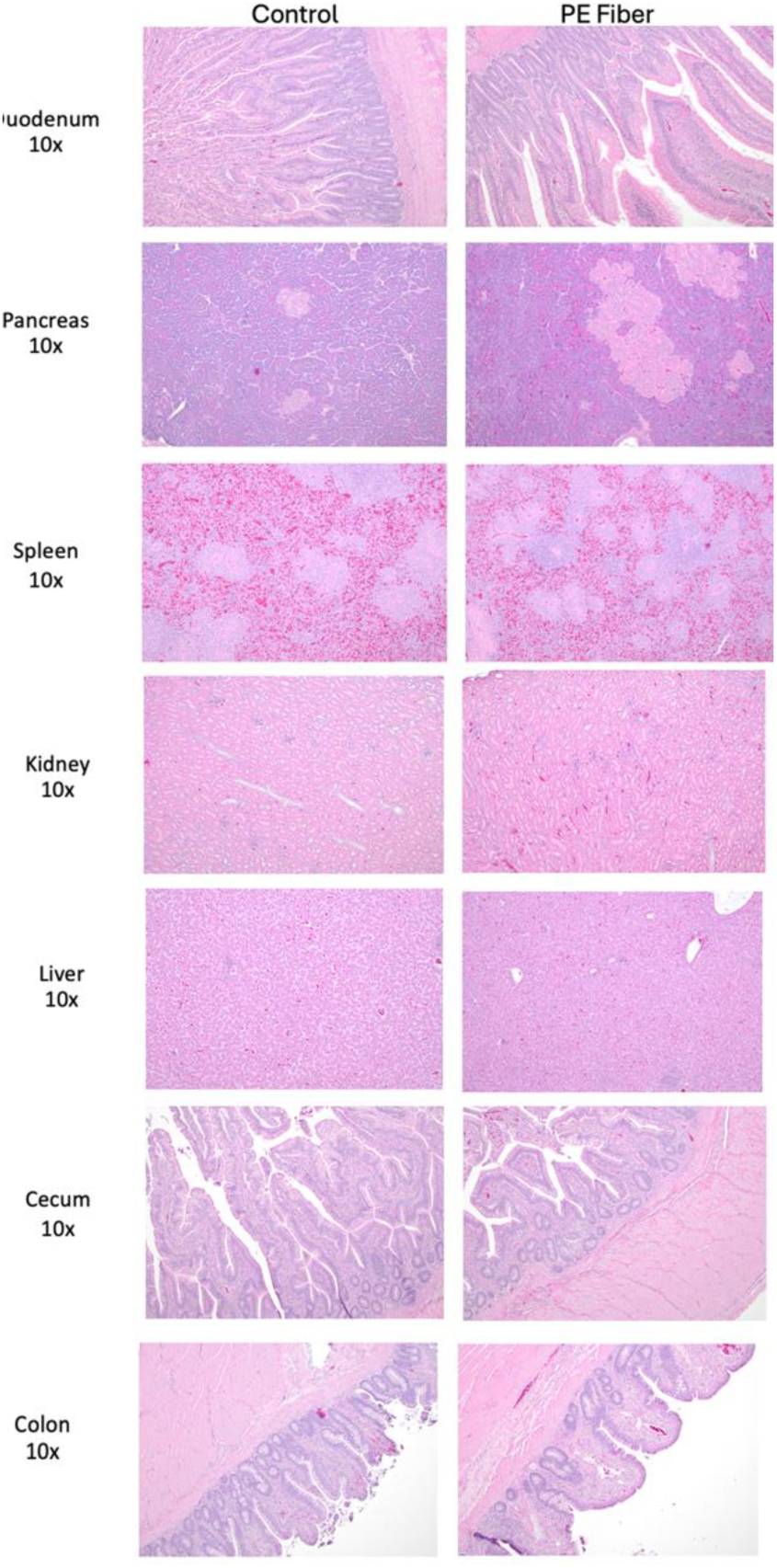
Hematoxylin and eosin staining of control and +PE fiber broilers. Sections included duodenum, pancreas, kidney, spleen, cecum, liver and colon. The images are representative of each treatment group.

**Figure S4.**
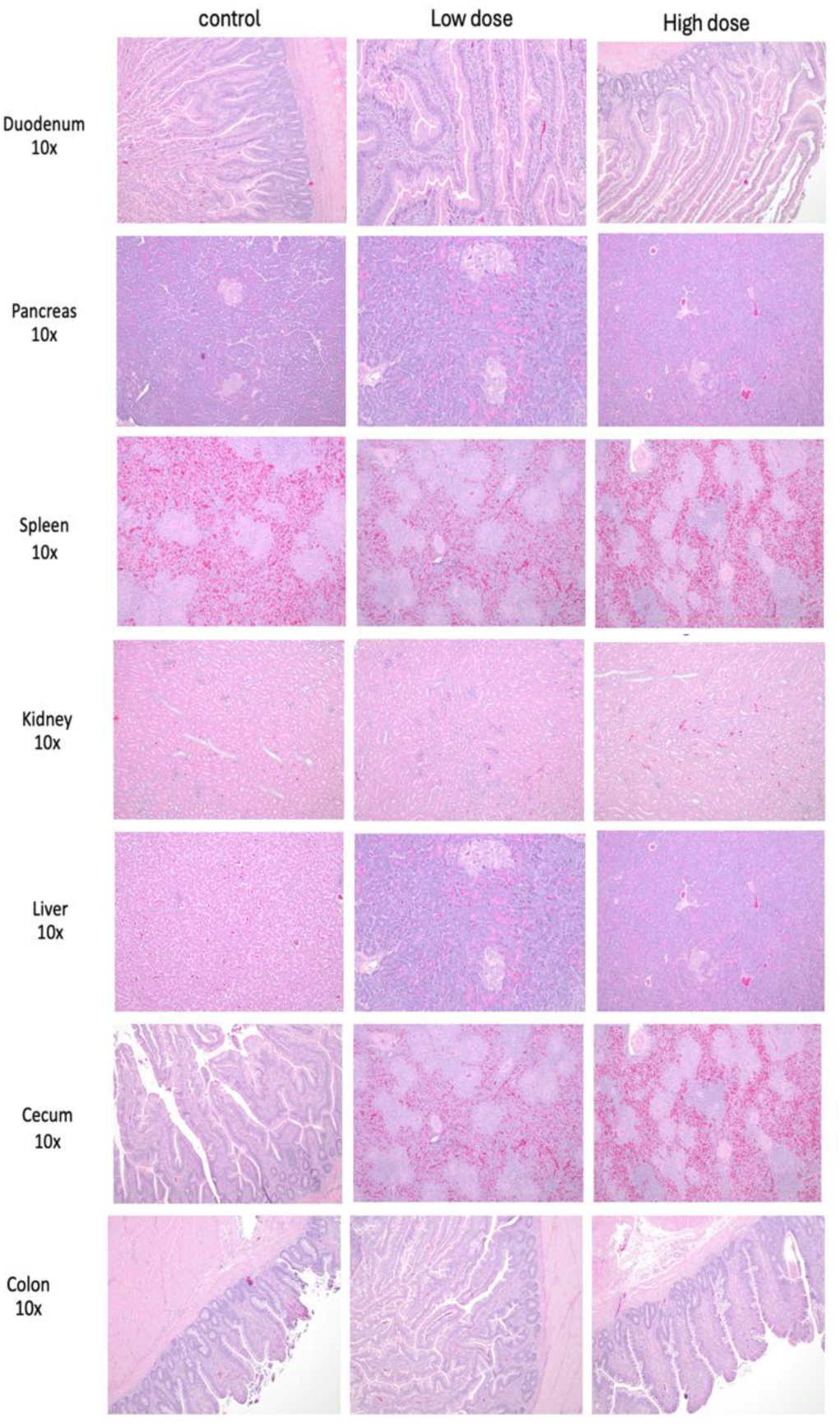
Hematoxylin and eosin staining of control. Low dose and high dose broilers. Sections included duodenum, pancreas, spleen, kidney, spleen, cecum, liver and colon. The images are representative of each treatment group.

**Figure S5.**
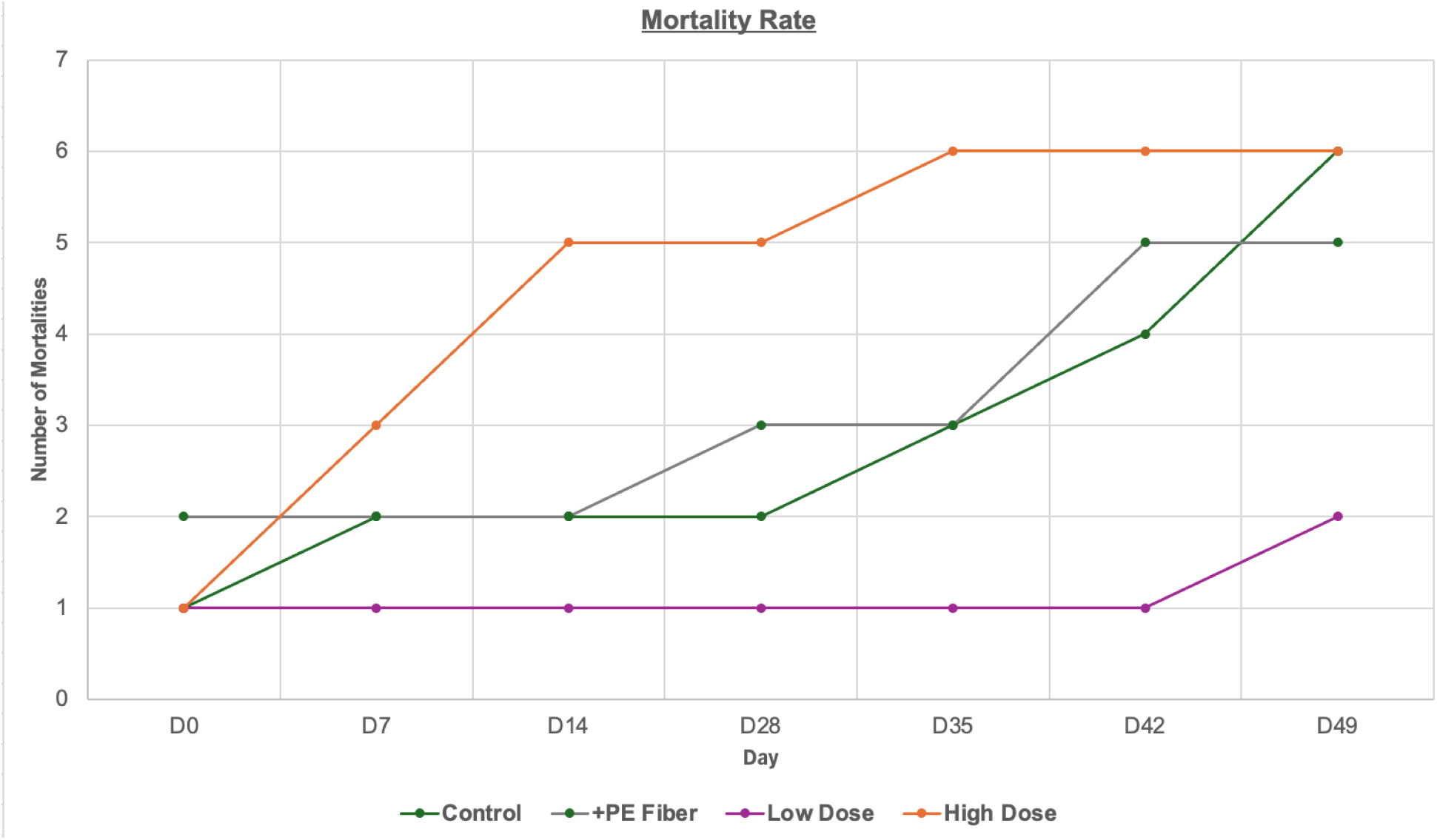
Mortality rate from the acclimation period (D0) through study termination (D49) for control (green), +PE Fiber(grey), low dose (purple) and high dose(orange).

**Figure S6.**
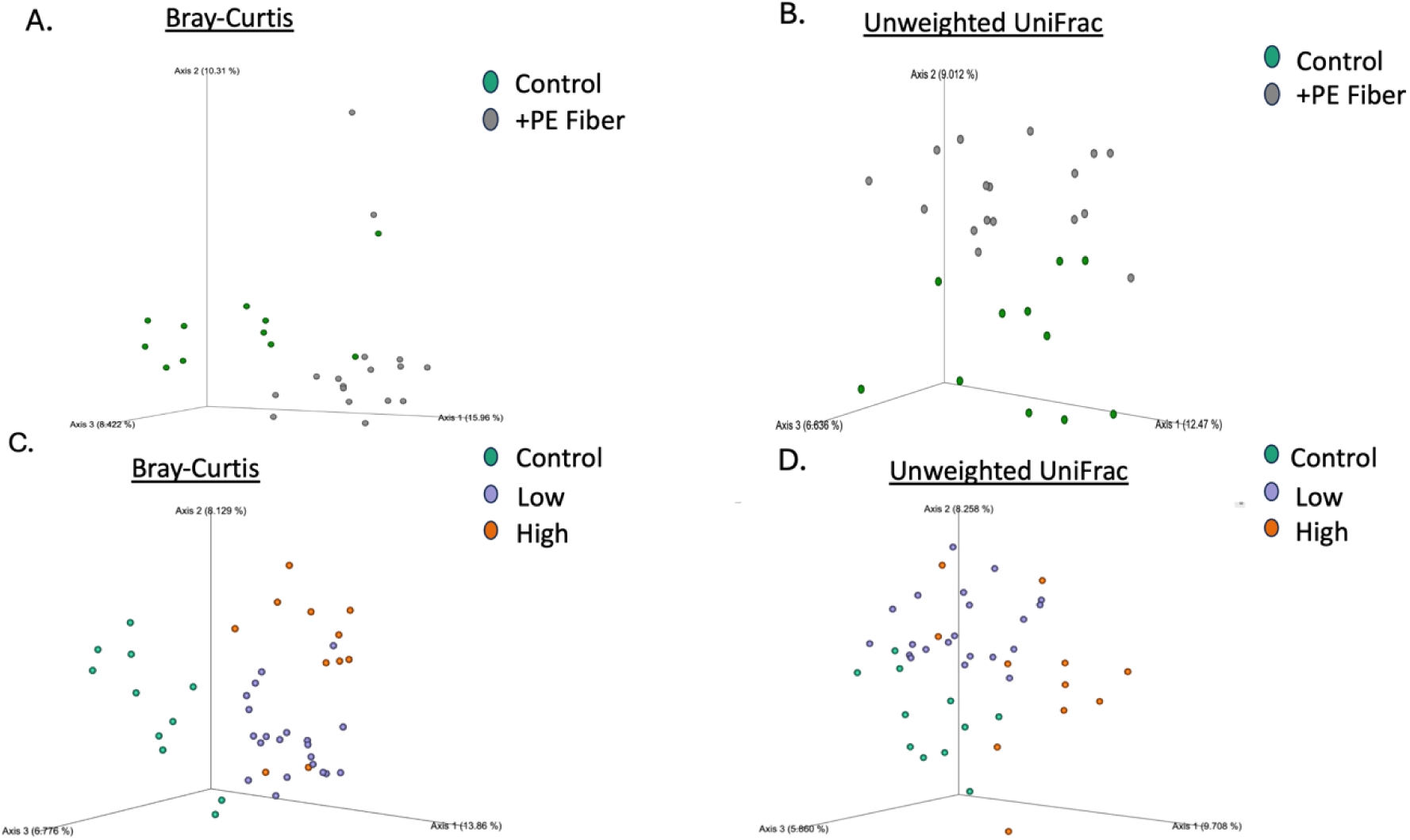
Principal component analysis for A) Bray-Curtis analysis comparing control and +PE Fiber treatment groups, B) Weighted UniFrac analysis control and +PE Fiber treatment groups, C) Bray-Curtis analysis comparing control, low dose and high dose treatment groups, and D) Weighted UniFrac analysis control, low dose and high dose treatment groups.

**Table S1.** Guaranteed analysis of poultry feed.^1^.

| Ingredients | % |
| --- | --- |
| Crude protein | 21 |
| Lysine, minimum | 1.0 |
| Methionine, minimum | 0.42 |
| Crude Fat, minimum | 4.0 |
| Crude Fiber, maximum | 6.0 |
| Calcium, minimum | 1.0 |
| Calcium, maximum | 1.2 |
| Phosphorus, minimum | 0.55 |
| Salt, minimum | 0.15 |
| Salt, maximum | 0.5 |
| Sodium, minimum | 0.15 |
| Sodium, maximum | 0.3 |

<sup>1</sup>.Agrimaster 21% Meat Producer Poultry Feed Crumbles; Blain # 334839, Mfr # 3931

**Table S2.** Ingredients list of poultry feed.^1^.

| Ingredients | %<sup>2</sup> |
| --- | --- |
| Processed Grain By-Products | -- |
| Grain Products Dehulled Soybean Meal | -- |
| Canola Meal | -- |
| Animal Protein Products | -- |
| Calcium Carbonate | -- |
| Salt | -- |
| DL-Methionine | -- |
| Yeast Culture | -- |
| L-Lysine | -- |
| Propionic Acid (a preservative) | -- |
| Dried Aspergillus niger Fermentation Extract | -- |
| Vitamin A Supplement | -- |
| Vitamin D3 Supplement | -- |
| Vitamin E Supplement | -- |
| Vitamin B12 Supplement | -- |
| Niacin Supplement | -- |
| d-Calcium Pantothenate | -- |
| Folic Acid | -- |
| Menadione Sodium Bisulfite Complex (source of Vitamin K activity) | -- |
| Riboflavin Supplement | -- |
| Pyridoxine Hydrochloride | -- |
| Thiamine Mononitrate | -- |
| Biotin | -- |
| Manganous Oxide | -- |
| Manganese Sulfate | -- |
| Ferrous Sulfate | -- |
| Copper Chloride | -- |
| Copper Sulfate | -- |
| Zinc Oxide | -- |
| Zinc Sulfate | -- |
| Ethylenediamine Dihydroiodide | -- |
| Sodium Selenite | -- |
| Dried Bifidobacterium thermophilum Fermentation Product | -- |
| Dried Enterococcus faecium Fermentation Product | -- |
| Dried Lactobacillus acidophilus Fermentation Product | -- |
| Dried Lactobacillus casei Fermentation Product. | -- |
| 1. AgriMaster 21% Meat Producer Poultry Feed Crumbles; Blain # 334839, Mfr # 3931 |  |
| 2. Percentages not available for ingredients list. |  |

**Table S3.** Kruskal-Wallis pairwise analysis results for α-diversity metrics comparing control and +PE Fiber treatment groups.

| Table S3. Kruskal-Wallis pairwise analysis results for α-diversity metrics comparing control and +PE Fiber treatment groups. | | | | |
| --- | --- | --- | --- | --- |
| Group 1 | Group 2 | α-diversity metric | | |
|  |  | Shannon's Diversity Index |  |  |
| +PE Fiber (n=17) | Control (n=11) | H | P-Value | Q-Value |
|  |  | 2.95 | 0.09 | 0.09 |
|  |  | Pielou's Evenness |  |  |
| +PE Fiber (n=17) | Control (n =11) | H | P-Value | Q-Value |
|  |  | 0.93 | 0.33 | 0.33 |
|  |  | Faith's phylogenetic Diversity |  |  |
| +PE Fiber (n=17) | Control (n=11) | H | P-Value | Q-Value |
|  |  | 2.63 | 0.1 | 0.1 |
|  |  | Observed Features |  |  |
| +PE Fiber (n=17) | Control (n=11) | H | P-Value | Q-Value |
|  |  | 2.2 | 0.14 | 0.14 |

**Table S4.** Analysis of control (n=11) and +PE Fiber (n=17) treatment effects based on β-diversity metrics as determined with PERMANOVA (P-value ≤ 0.05; Q-value ≤ 0.05).

| Table S4. Analysis of control (n=11) and +PE Fiber (n=17) treatment effects based on β-diversity metrics as determined with PERMANOVA (P-value ≤ 0.05; Q-value ≤ 0.05).<sup>a</sup> | | | | | |
| --- | --- | --- | --- | --- | --- |
| Metric | Test statistic | Sample size | Test statistic | P-value | Q-value |
| Bray-Curtis | Pseudo-F | 28 | 4.09 | 0.00 | 0.00 |
| Jaccard | Pseudo-F | 28 | 2.12 | 0.00 | 0.00 |
| Weighted UniFrac | Pseudo-F | 28 | 2.21 | 0.00 | 0.00 |
| Unweighted UniFrac | Pseudo-F | 28 | 2.16 | 0.00 | 0.00 |
| <sup>a</sup> PERMANOVA performed with 999 permutations. |  |  |  |  |  |

**Table S5.** Analysis of control (n=11), low dose (n=21) and high dose (n=11) treatment effects based on α-diversity metrics as determined with ANOVA.

| Table S5. Analysis of control (n=11), low dose (n=21) and high dose (n=11) treatment effects based on α-diversity metrics as determined with ANOVA. | | | | | |
| --- | --- | --- | --- | --- | --- |
| Metric | Model | Sum_sq | Df | F | PR(>F) |
| Faith's Phylogenetic Diversity | Treatment | 43.79 | 2.0 | 10.48 | 0.00 |
|  | Residual | 83.57 | 40.0 |  |  |
| Shannon's entropy | Treatment | 0.84 | 2.0 | 3.48 | 0.04 |
|  | Residual | 4.84 | 40.0 |  |  |
| Pielou's evenness | Treatment | 0.00 | 2.0 | 1.32 | 0.29 |
|  | Residual | 0.04 | 40.0 |  |  |
| Observed Features | Treatment | 21,140.86 | 2.0 | 7.75 | 0.00 |
|  | Residual | 54,524.21 | 40.0 |  |  |

**Table S6.** Kruskal-Wallis pairwise analysis results for a-diversity metrics.

| Table S6. Kruskal-Wallis pairwise analysis results for α-diversity metrics. | | | | |
| --- | --- | --- | --- | --- |
| Group 1 | Group 2 | $\alpha$ -diversity metrics <sup>a</sup> | | |
|  |  | Shannon's Diversity Index |  |  |
|  |  | H | P-value | Q-value |
| High dose mixture (n =11) | Control (n= 11) | 2.59 | 0.11 | 0.16 |
| Low dose mixture (n =21) | Control (n= 11) | 0.66 | 0.41 | 0.41 |
| High dose mixture (n =11) | Low dose mixture (n= 21) | 6.14 | **0.01** | **0.04** |
|  |  | Pielou's Evenness |  |  |
|  |  | H | P-Value | Q-Value |
| High dose mixture (n =11) | Control (n= 11) | 1.99 | 0.16 | 0.41 |
| Low dose mixture (n =21) | Control (n= 11) | 0.33 | 0.56 | 0.56 |
| High dose mixture (n =11) | Low dose mixture (n= 21) | 1.19 | 0.26 | 0.41 |
|  |  | Faith's phylogenetic Diversity |  |  |
|  |  | H | P-Value | Q-Value |
| High dose mixture (n =11) | Control (n= 11) | 7.07 | **0.01** | **0.01** |
| Low dose mixture (n =21) | Control (n= 11) | 0.01 | 0.92 | 0.92 |
| High dose mixture (n =11) | Low dose mixture (n= 21) | 11.78 | **0.00** | **0.00** |
|  |  | Observed Features |  |  |
|  |  | H | P-Value | Q-Value |
| High dose mixture (n =11) | Control (n= 11) | 7.25 | **0.01** | **0.01** |
| Low dose mixture (n =21) | Control (n= 11) | 0.01 | 0.94 | 0.94 |
| High dose mixture (n =11) | Low dose mixture (n= 21) | 7.61 | **0.01** | **0.01** |
| <sup>a</sup>. Bolded values indicate significance (P-value ≤ 0.05; Q-value ≤ 0.05). |  |  |  |  |

**Table S7.**
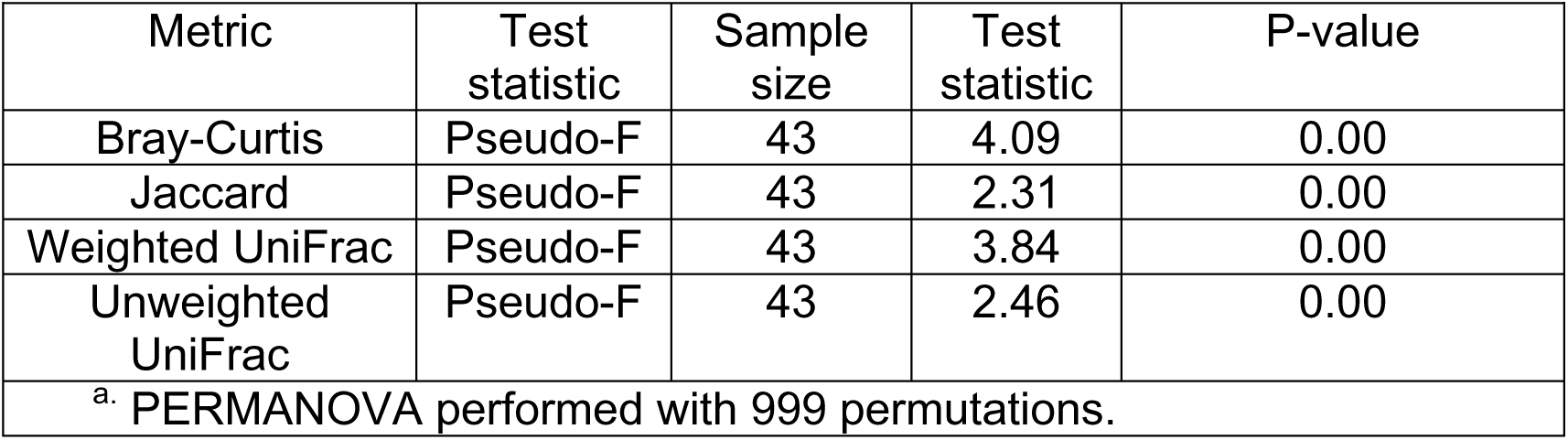
Analysis of control (n=11), low dose (n=21) and high dose (n=11) treatment effects based on β-diversity metrics as determined with PERMANOVA (P-value ≤ 0.05).^a^.

